# Amino acid substitutions hydrophilizing the core of titin domains cause dilated cardiomyopathy

**DOI:** 10.1101/2023.06.02.543467

**Authors:** Ines Martinez-Martin, Audrey Crousilles, Juan Pablo Ochoa, Diana Velazquez-Carreras, Simon A. Mortensen, Elias Herrero-Galan, Fernando Dominguez, Pablo Garcia-Pavia, David de Sancho, Matthias Wilmanns, Jorge Alegre-Cebollada

## Abstract

The underlying genetic defect in most cases of dilated cardiomyopathy (DCM), a common inherited heart disease, remains unknown. Intriguingly, many patients carry single missense variants of uncertain pathogenicity targeting the giant protein titin, a fundamental sarcomere component. To explore the deleterious potential of these variants, we first solved the wild-type and mutant crystal structures of I21, the titin domain targeted by pathogenic variant p.C3575S. Although both structures are remarkably similar, the increase in hydrophilicity of deeply buried position 3575 strongly destabilizes the mutant domain, a scenario supported by molecular dynamics simulations and by biochemical assays that show no disulfide involving C3575. Prompted by these observations, we have found that thousands of similar hydrophilizing variants associate specifically with DCM. Hence, our results imply that titin domain destabilization causes DCM, a conceptual framework that not only informs pathogenicity assessment of gene variants but also points to therapeutic strategies counterbalancing protein destabilization.

## Introduction

Dilated cardiomyopathy (DCM) is a cardiac disease characterized by left ventricle dilatation and systolic dysfunction that cannot be explained solely by abnormal loading conditions or coronary artery disease. Common complications include arrhythmias and heart failure^1^, making DCM the most frequent cause of heart transplantation^2^. In the last two decades, clinical management of DCM patients and their families has benefited from identification of causative genetic variants^3^. For instance, non-carrier relatives can be assured they are not at higher risk of developing DCM than the general population, which has a profound impact in their life planning^4^. However, at the present time the genetic origin of DCM cannot be identified in ∼60-70% of families, who therefore cannot take advantage of genetic-based improved care. Instead, genetic analysis of these otherwise genotype-negative individuals typically detects variants for which there is not enough evidence to support pathogenicity, the so-called variants of uncertain significance (VUS)^5^. VUS have no use in the clinic; hence, there is a pressing need for strategies able to discriminate pathogenic variants from clinically innocuous, benign polymorphisms.

Truncating variants in the titin gene (*TTN*) leading to titin loss of function are a major cause of familial DCM, accounting for approximately 25% of cases^6,7^. Titin is a structural protein that bridges the Z-disk and M-line of sarcomeres in myocytes acting as a molecular spring and scaffold for a variety of protein interactions^8,9^. The elastic I-band part of titin is composed of serially linked immunoglobulin-like (Ig) domains and random coil regions, whereas the rigid A-band contains both Ig and fibronectin type III (FnIII) domains (**Figure 1A**)^10^. Titin is regulated by alternative splicing, which gives rise to isoforms of different length that modulate myocyte stiffness^11^. The long and compliant N2BA and the shorter and more rigid N2B are the two major isoforms in the adult human heart, in which they are present at a 30:70 proportion, respectively^8^.

**Figure 1.**
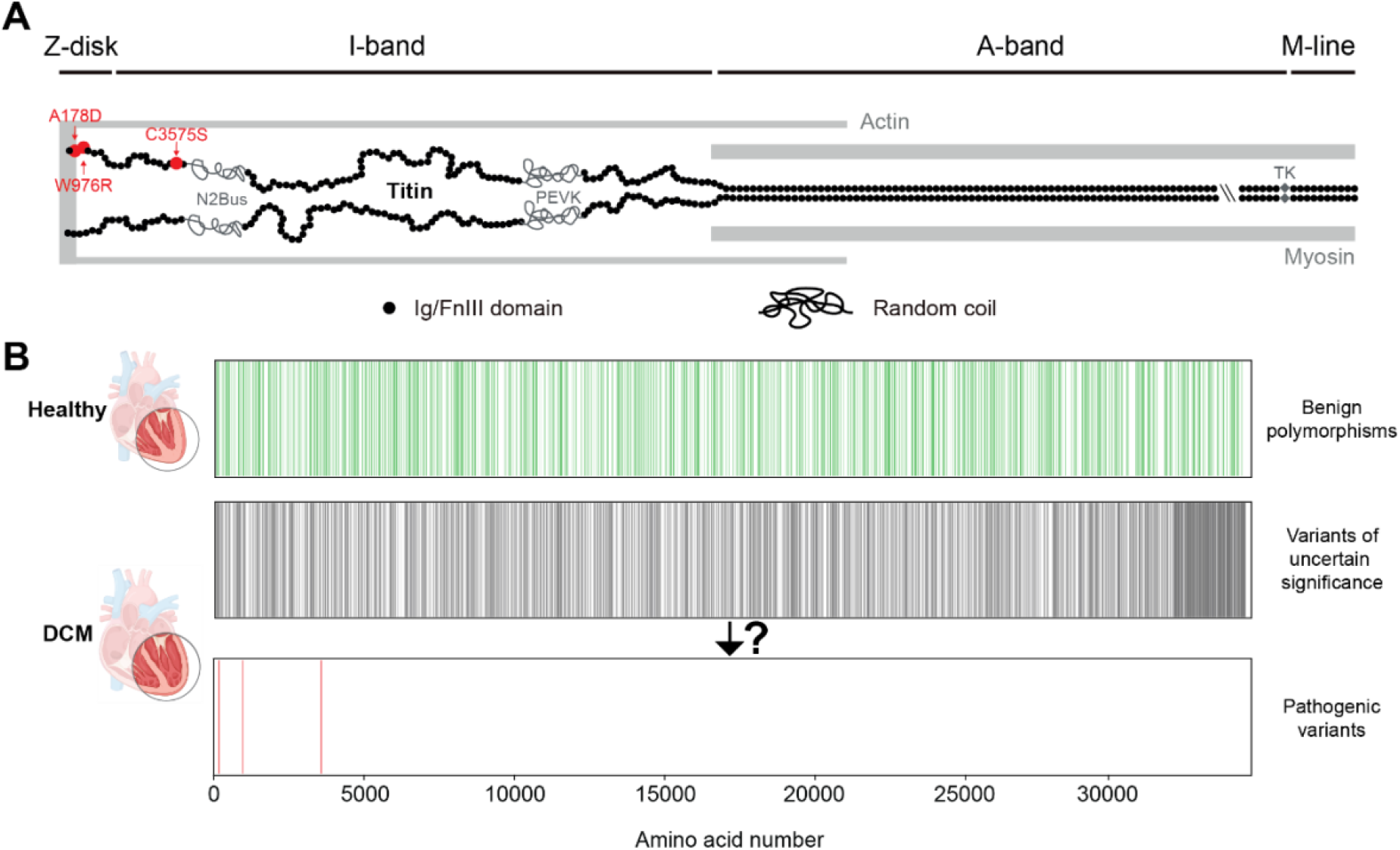
Landscape of titin missense variants in DCM: **A.** Representation of one half a sarcomere (not to scale). N2BA titin molecules are colored in black. Ig and FnIII domains are represented as circles. Positions of PEVK and N2Bus random coil regions are shown. The gap indicated by double backslash corresponds to 100 domains in the A-band that are not represented for simplicity. To date, only three missense variants in titin have been demonstrated to cause DCM (shown in red). **B.** Location of missense variants in the sequence of titin. Benign polymorphisms are represented in green (top), while variants of uncertain significance identified in DCM patients are colored in grey (middle). Pinpointing pathogenic variants among these VUS has not been possible so far. In red (bottom), location of the three pathogenic variants shown in panel A.

Remarkably, missense variants in titin are found frequently in DCM individuals^6,12^. However, in the absence of further information, whether these variants are pathogenic or not is impossible to ascertain since benign polymorphisms in titin are also common in the general population^13^. As a consequence, most titin missense variants have remained classified as VUS^12,14^. Indeed, only three titin missense variants (p.A178D, p.W976R and p.C3575S) have been linked to DCM so far based on robust cosegregation with disease^15–19^(**Figure 1**). It is tempting to hypothesize that the pathogenic features of these DCM-causing variants could also be present in some of the myriad other missense VUS in titin, potentially enabling their reclassification as pathogenic. In this regard, it is interesting to note that the three aforementioned DCM-causing missense variants destabilize their parent domains^15,17,19^. Hence, we hypothesized that uncovering molecular mechanisms inducing titin domain destabilization could lead to identification of DCM-causing missense variants.

To examine this hypothesis, here we have studied *TTN* p.C3575S (reference Uniprot sequence Q8WZ42-1), the titin missense variant with the strongest association with DCM so far^19^. Using X-ray crystallography, protein biochemistry, molecular simulations and bioinformatics analyses, we show that the strong destabilization induced by the cysteine to serine substitution results from a substantial decrease in hydrophobicity of the core of the parent I21 domain. Building on these results, we have found that similar hydrophilizing variants targeting titin domains are enriched in DCM cohorts, an observation that places these variants as a causative factor of DCM.

## Results

### Crystal structure of wild-type and C3575S I21 titin domains

We first solved the structure of wild-type (WT) I21 domain to 1.95Å resolution (PDB ID 8OVU, diffraction data statistics and model refinement parameters are in **Supplementary Table S1**). The asymmetric unit of the crystal contains two copies of the domain that show highly similar structures (root mean square displacement, RMSD, is 0.50Å, **Figure 2A, Supplementary Figure S1A**). The fold of the I21 domain falls within the intermediate set (type-I) of the immunoglobulin superfamily^20,21^. Two antiparallel β-sheets formed by four beta strands (A-B-E-D and A’-G-F-C) are packed in a β-sandwich. Residue C3575 is located on strand F and is deeply buried in the core of the domain (solvent accessible surface (SAS) < 0.01 Å^2^). In one of the chains, a Mg^2+^ atom is coordinated in octahedral geometry with D3548 and five water molecules (**Figure 2B**). Mg^2+^ binding is accompanied by minor rearrangement of surrounding side chains, including rotation of D3548 and E3568 (**Figure 2C**).

**Figure 2.**
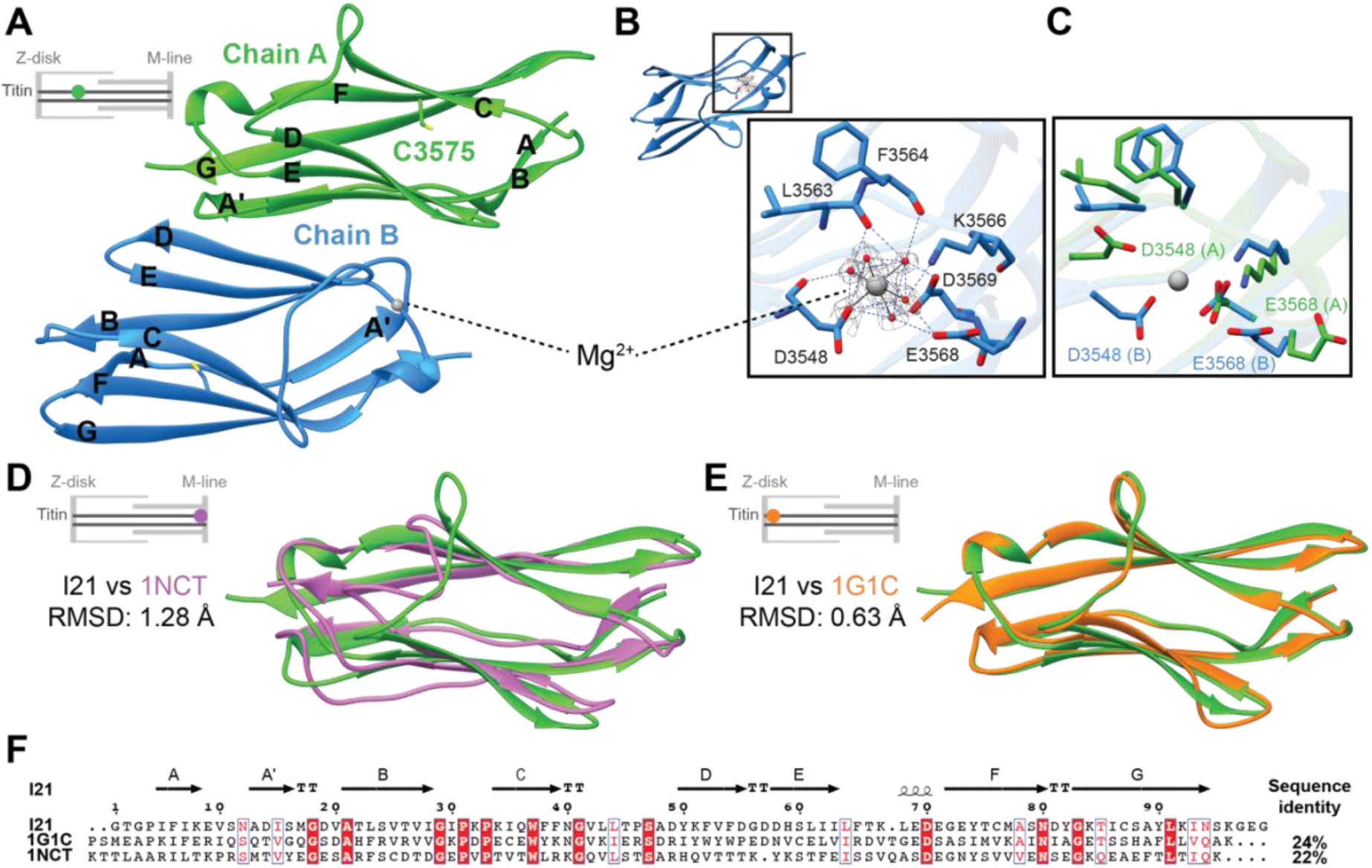
The I21 domain of titin shows a typical Ig fold. **A**. Ribbon cartoon of the asymmetric unit of the crystal containing the I21 domain. C3575 is shown as sticks with sulfur atoms in yellow. Chain B (blue) interacts with a Mg^2+^ atom depicted as a gray sphere. **B**. Mg^2+^-binding site in the crystal structure of I21. Residues and water molecules (red spheres) coordinating the metal are shown (interactions as dashed lines). The electron density surrounding the Mg^2+^ is contoured at 1.2σ (gray mesh). **C**. Structural alignment of the Mg^2+^-binding site in chains A (green) and B (blue). **D**. Ribbon cartoon of the structural alignment of domains I21 and 1NCT. **E**. Ribbon cartoon of the structural alignment of domains I21 and 1G1C. **F**. Sequence alignment of I21, 1G1C and 1NCT domains. The secondary structure assignment of I21 is shown on top. Sequence identities between I21 and 1G1C and 1NCT appear on the right. Insets of panels A, D and E indicate the location of the corresponding titin domains.

To examine whether I21 has the typical titin Ig domain fold, we aligned I21 to 13 available high-resolution structures of Ig domains of human titin covering all relevant titin regions (**Supplementary Table S2**). We found that all domains show remarkably high structural similarity. The most dissimilar domain is 1NCT, located in the M-line (RMSD 1.28 Å, **Figure 2D**), whereas the I-band domain 1G1C shows the lowest RMSD with I21 (0.63 Å, **Figure 2E**). This high structural similarity is not strongly correlated with sequence identity (**Figure 2F**), as observed in many other members of the Ig superfamily^22^.

Next, we solved the crystal structure of the I21 C3575S mutant domain to 2.2 Å resolution (PDB ID 8P35, diffraction data statistics and model refinement parameters in **Supplementary Table S1**). The asymmetric unit of the crystal contains six copies of the domain (RMSD range: 0.25-0.48 Å using chain A as a reference, **Figure 3A**). Strikingly, despite the highly destabilizing effect of C3575S^19^, the mutation does not induce any noticeable change in the general fold of the domain (RMSD I21 WT vs I21 C3575S = 0.54 Å, **Figure 3B**). Indeed, the side chains of 3575 and surrounding residues align to a great extent in both structures. There are only slight rotations of I3534 and V3509, which however are unlikely to affect the main interactions of the fold (**Figure 3C**).

**Figure 3.**
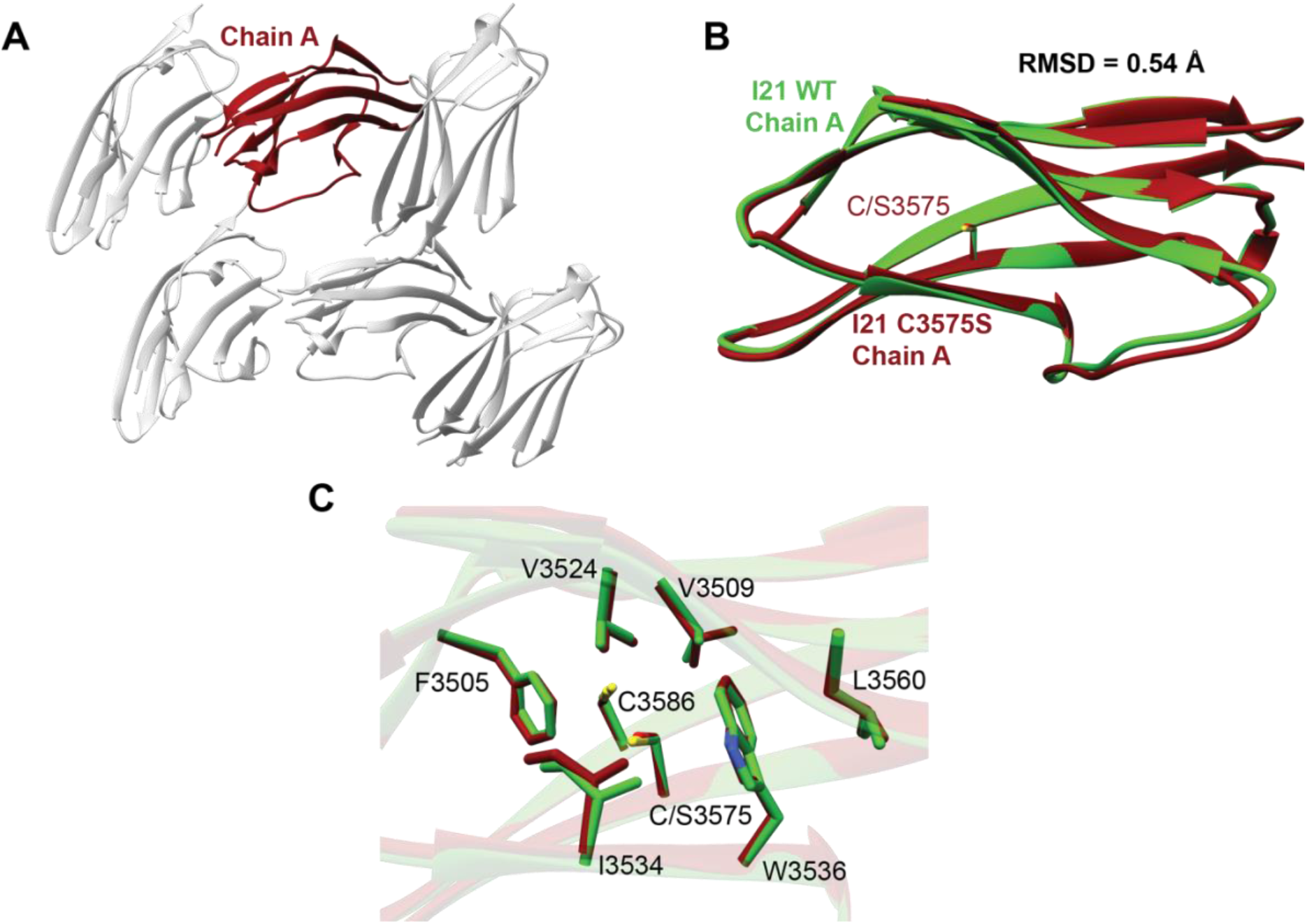
The C3575S mutation does not induce major structural changes in domain I21. **A**. Ribbon cartoon of the asymmetric unit of the crystal containing the I21 C3575S domain. Chain A is colored in dark red, and chains B-F in light gray. **B**. Structural alignment of I21 WT (green) and C3575S (dark red). The RMSD of the alignment is 0.54 Å. Position 3575 is represented with sticks. **C**. Detailed view of the region surrounding position 3575 in the structural alignment of I21 WT (green) and I21 C3575S (dark red).

### I21 C3575 domain destabilization is not caused by loss of a disulfide bond

The structure of the I21 WT domain shows a second cysteine residue, C3586, at a distance from C3575 that is compatible with disulfide bond formation (**Figure 4A**). The equivalent positions in many titin Ig domains are typically occupied by cysteines that have been shown to be able to form disulfide bonds^23,24^. The destabilizing effect of the C3575S mutation could stem from the inability of serine to establish a disulfide bond with C3586. However, we found no evidence of disulfide bond formation in the experimental electron density of the WT domain (**Supplementary Figure S1B**). To confirm that I21 domain destabilization is not caused by loss of disulfide bond C3575-C3586, we examined the stability of WT and mutant domain preparations whose redox state was evaluated concomitantly using a biochemical NEM-PEGylation assay. In this assay, both domains were incubated with an N-ethylmaleimide-functionalized PEG molecule that binds covalently to reduced cysteines, causing a mobility shift in SDS-PAGE electrophoresis (**Figure 4B**). As expected, NEM-PEG-treated I21 C3575S domain shows a shift corresponding to the addition of a single PEG molecule (**Figure 4C**). The WT domain also shows full modification of cysteines, but in this case the shift in electrophoretic mobility corresponds to 2 PEG molecules (**Figure 4C)**. Hence, results of the NEM-PEGylation assay demonstrate that both domain preparations did not contain disulfide bonds nor any other form of cysteine oxidation. Next, using circular dichroism (CD) with these fully reduced protein preparations, we verified that C3575S causes strong destabilization of I21. Specifically, the WT domain has a melting temperature (T_m_) of 52 ± 0.3 ºC, while the T_m_ of mutant I21 is 38 ± 0.5 ºC (errors are SD from the sigmoidal fit; **Figure 4D**). Hence, our results confirm that the loss of thermal stability of I21 C3575 is not caused by the loss of a disulfide bond.

**Figure 4.**
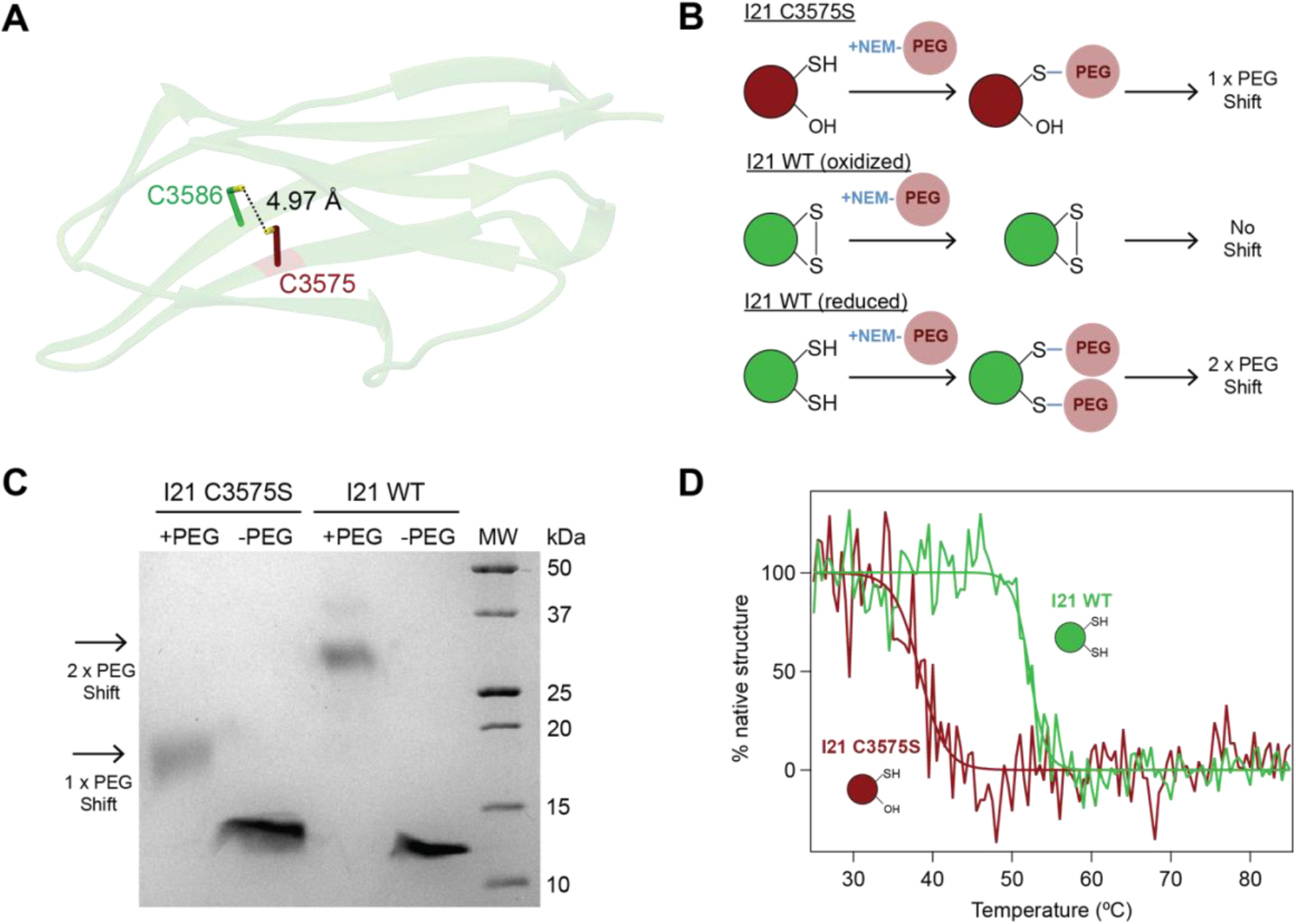
I21 domain destabilization induced by C3575S is not related to the disruption of a disulfide bond. **A**. Cartoon representation of I21 WT depicting the two cysteines of the domain as sticks with sulfur atoms in yellow. The dashed line indicates the distance between sulfur atoms (4.97 Å). **B**. Schematic representation of the NEM-PEG assay applied to the I21 C3575S (dark red) and WT (green) domains. **C**. 17% SDS-PAGE showing the results of the NEM-PEG assay. MW: molecular weight markers (Biorad Precision Plus Protein) **D**. Thermal unfolding curves of I21 WT (green) and C3575S (dark red) measured by tracking CD signal at 215 nm. Sigmoidal fits to the data are shown.

### C3575S perturbs the I21 equilibrium folding free energy

To characterize the thermodynamic changes induced by the C3575S mutation, we estimated the associated variation in folding free energy (ΔΔG) from molecular dynamics (MD) simulations using an alchemical formalism. This approach, which has been applied to calculate the effects of mutations in protein stability^25^ and ligand binding affinity^26^, is based on a thermodynamic cycle that includes both physical (un)folding reactions and unphysical mutation transitions (**Figure 5A**). According to this cycle, ΔΔG can be obtained from the difference in ΔG of the mutation transitions in the unfolded and folded states (ΔG_m,unfolded_ and ΔG_m,folded_, respectively). This approach greatly alleviates the computational expense of the free energy calculation compared to the simulation of reversible folding and unfolding events for the two forms of the protein. ΔG_m_ values are calculated from atomistic, explicit-solvent MD trajectories in which the residue of interest is artificially switched from WT to mutant and vice versa (**Figure 5A,B**). Using two different MD force fields, our alchemical calculations indicate that mutation C3575S induces high thermodynamic destabilization (average ΔΔG = 16.5 ± 0.5 kJ/mol) (**Figure 5C**), which correlates well with the 14 ºC decrease in *T*_*m*_ we have measured experimentally ^27^.

**Figure 5.**
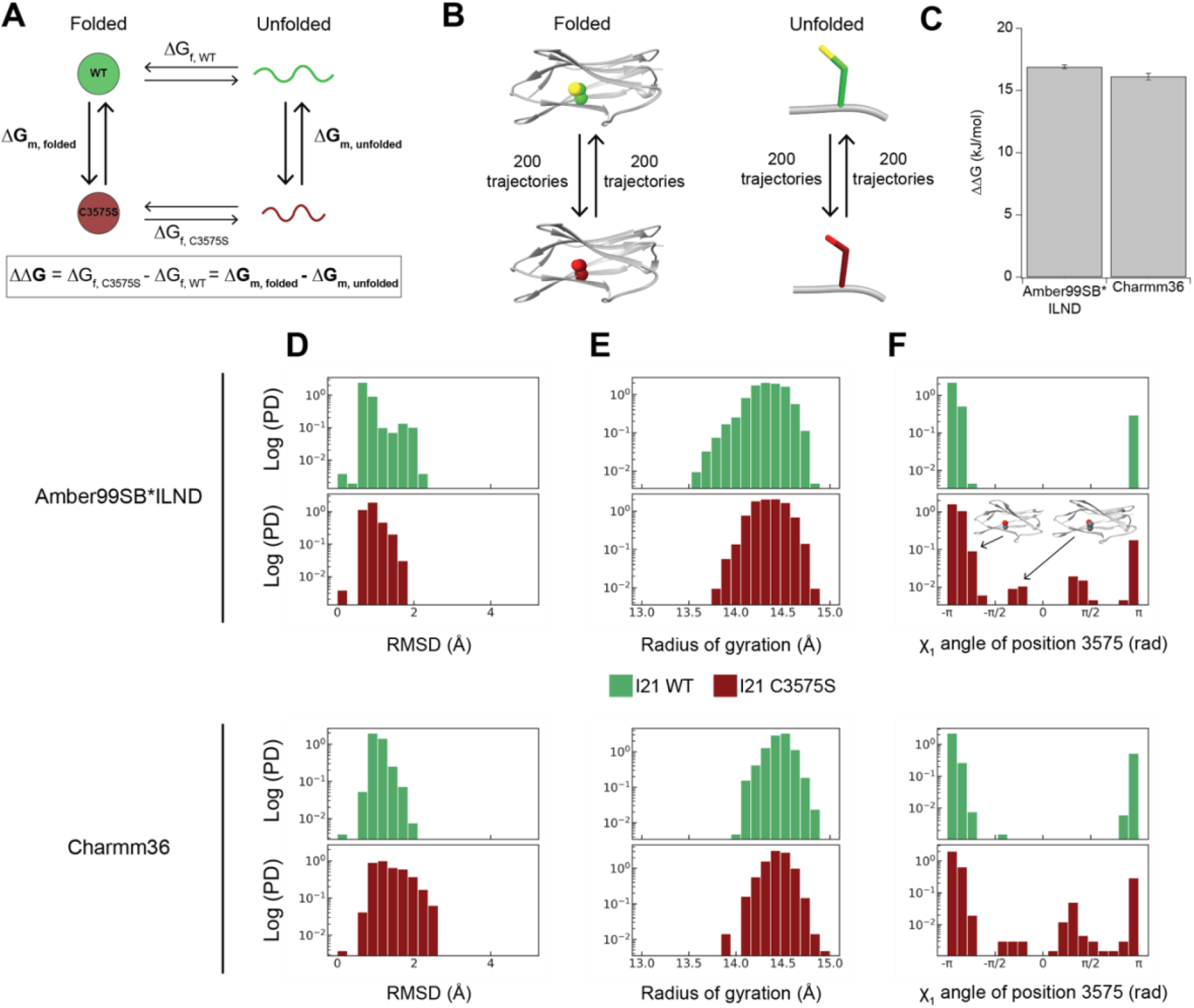
Molecular dynamics simulations capture destabilization of I21 C3575S domain. **A.** Thermodynamic cycle used for ΔΔG calculations. **B**. Non-physical transitions used by the alchemical MD-based free-energy calculation. **C**. ΔΔG values using two different MD force fields. Error bars show standard deviation. **D**. Probability density histograms of global RMSD values (alpha carbons) in the folded state of wild-type and mutant domains. **E**. Probability density histograms of radius of gyration in the folded state of wild-type and mutant domains. **F**. Probability density histograms of X1 angle values of position 3575 in the folded state of wild-type and mutant domains. A ribbon cartoon of the I21 domain illustrates the X1 angles seen in the trajectories. Histograms in panels D-F were calculated pooling data from 2 independent trajectories with Amber99SB*ILND (top) or Charmm36 (bottom) force fields.

To look for insights into the molecular mechanisms of destabilization, we analyzed the long equilibrium MD trajectories used to seed the switching simulations. Despite the remarkable destabilization induced by C3575S, we could not detect any noticeable effect of the mutation on the alpha carbon RMSD distributions, on the global conformational mobility of the folded domain or on its radius of gyration (**Figure 5D,E, Supplementary Figure S2A-F**). Although residue 3575 remained buried throughout the simulated time for both WT and mutant (**Supplementary Figure S2G-H)**, the serine in the C3575S folded state explored more frequently alternative rotameric states characterized by X_1_ angles between -π/2 and π/2 (**Figure 5F, Supplementary Figures S2I-J**). This higher conformational heterogeneity suggests entropic stabilization of the native state of the C3575S domain^28^. Using the Shannon entropy formula, we have estimated that the associated entropic stabilization of the mutant is 0.6-0.8 kJ/mol. Hence, this effect is small and easily compensated by the enthalpic stabilization of the Cys residue in the core of the folded domain. Indeed, according to statistical inter-residue contact energies for proteins^29^, the Cys to Ser substitution is expected to result in a 4-7 kJ/mol decrease per contact with nearby residues F3505, V3524, I3534, W3536, and C3586 (**Figure 3C**).

### Hydrophobicity loss in DCM-associated titin variants

Our MD simulations confirm the strong thermodynamic destabilization of the C3575S domain, which most likely originates from the loss of hydrophobic contacts induced by the cysteine to serine substitution. Indeed, most hydrophobicity scales, including the commonly used Kyte-Doolittle scale, agree that the hydrophobicity of cysteine is higher than that of serine^30,31^ (**Figure 6A**). Considering also that 3575 is the least solvent-accessible position in I21 (**Figure 6B**), it appears that C3575 is a fundamental residue to maintain key hydrophobic interactions in the core of the domain^32^. Accordingly, 99% of titin domains contain hydrophobic residues in the position equivalent to 3575 (**Figure 6C**), while only 55% residues are hydrophobic in the less buried position corresponding to vicinal C3586 (SAS = 10.63 Å^2^, **Figure 6B**,**D**).

**Figure 6.**
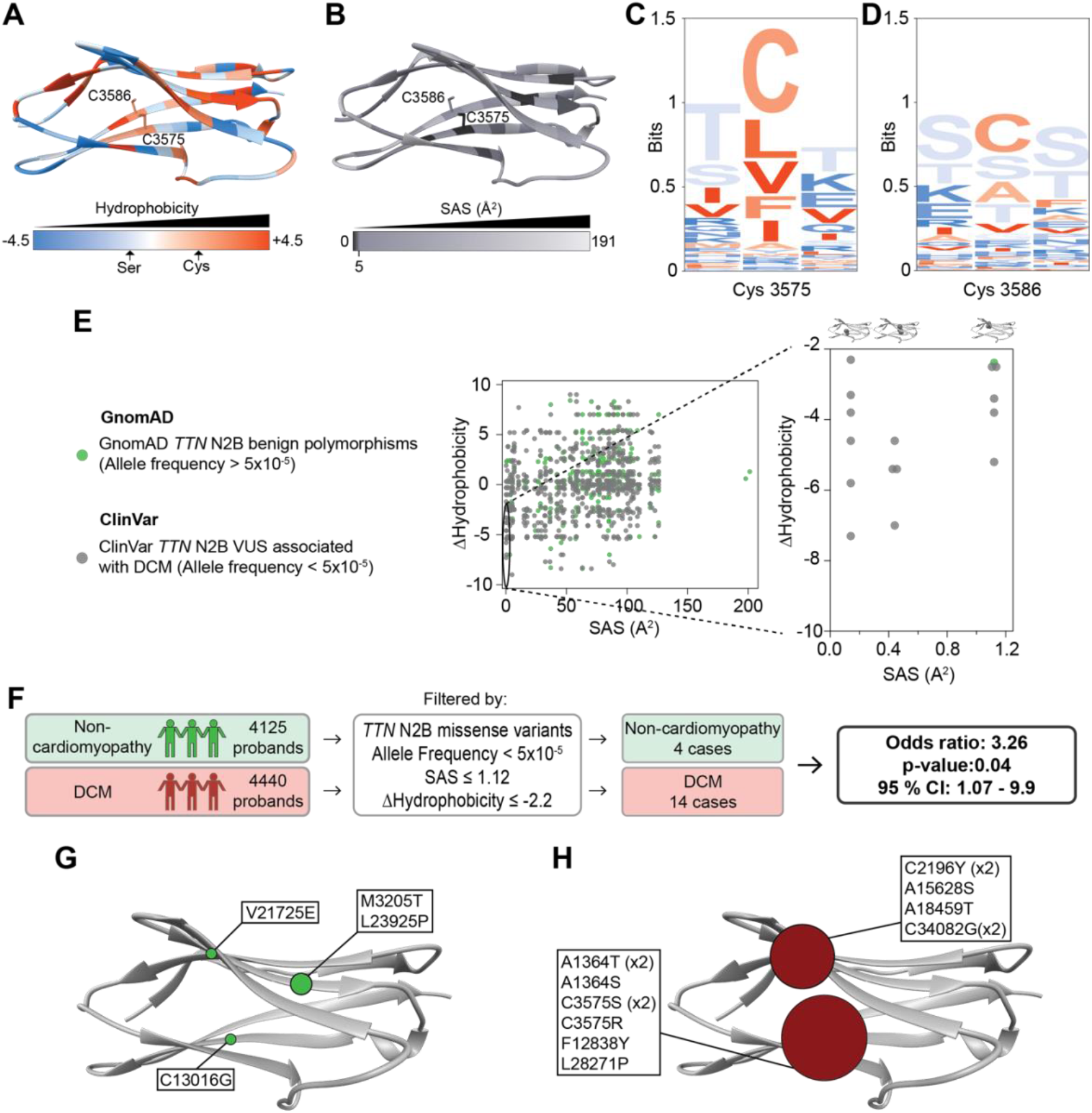
Variants hydrophilizing the core of titin Ig domains associate specifically with dilated cardiomyopathy. **A**. Cartoon ribbon representation of I21 WT colored according to residue hydrophobicity (Kyte-Doolittle scale). C3575 and C3586 are represented as sticks. **B**. Cartoon ribbon representation of I21 WT colored according to residue SAS. C3575 and C3586 are represented as sticks. **C, D**. Sequence logos of positions equivalent to 3574-3576 (C) and 3585-3587 (D) in the alignment of titin Ig domains^23^. Residues are colored according to their hydrophobicity. **E**. SAS and ΔHydrophobicity values of all DCM-associated VUS (grey) and benign polymorphisms (green). Zoom of variants with SAS < 1.12 Å^2^ and ΔHydrophobicity < -2.2 is on the right. **F**. Scheme of the case-control study showing specific association of hydrophilizing variants in the core of titin Ig domains with DCM. **G, H**. Positions of the hydrophilizing titin variants affecting deeply buried residues found in the non-cardiomyopathy (G, green) and DCM (H, dark red) cohorts. The size of the circles represents the number of variants found for each position.

To investigate if similar hydrophilizing variants cause DCM as p.C3575S, we examined whether they are specifically found in DCM populations. To this aim, we first calculated the changes in hydrophobicity (ΔHydrophobicity) according to the of thousands of publicly available VUS in ClinVar potentially associated with DCM that target Ig domains constitutively expressed in the N2B isoform of cardiac titin and, therefore, also present in the longer N2BA isoform. As controls, we used polymorphisms from gnomAD with allelic frequencies that are not compatible with the prevalence of DCM^33,34^. We plotted ΔHydrophobicity values according to the average SAS of the mutated position in titin Ig domains (**Supplementary Figure S3**). As expected, global results do not show noticeable differences in the distribution of DCM-associated VUS and benign polymorphisms. However, variants causing substantial drops in hydrophobicity (ΔHydrophobicity < -2.2) and targeting extremely buried positions (SAS < 1.12 Å^2^) are almost exclusively populated by DCM-associated VUS (p = 0.008, hypergeometric test) (**Figure 6E**).

Next, we examined to what extent variants showing those physicochemical features are enriched in DCM patients. To this aim, we performed an independent case-control study using a cohort of 4440 DCM cases with an unidentified genetic cause following evaluation with a 121 gene-next-generation-sequencing DCM panel, and 4125 control individuals referred for genetic testing due to conditions not related to DCM. Results show that hydrophilizing titin variants affecting deeply buried residues associate specifically with DCM with an odds ratio of 3.26 (p = 0.04, 95% Confidence Interval = 1.07 – 9.9) (**Figure 6F-H, Supplementary Table S3**).

## Discussion

The identification of genetic variants causing DCM is revolutionizing the clinical management of patients and their families^3,5^. However, currently only ∼30% DCM patients can benefit from improved care, i.e. those carrying well-known pathogenic variants^35,36^. This situation highlights the pressing need to identify DCM-causing variants, which typically relies on genetic screening of very large families in the quest for cosegregating candidates^17,19^ or identification of specific enrichment in large DCM cohorts^6^. Here, we have followed a third, mixed approach that first identifies pathogenic features in well-established DCM-causing variants, which are then screened for in VUS found in DCM patients.

Loss-of-function truncations in the titin gene are a common cause of DCM^6,14^. Prompted by this evidence, we and others have speculated that a fraction of titin missense variants could also result in protein loss of function leading to DCM^12–14^. However, to date only three titin missense variants have been shown to cause DCM (**Figure 1**). Among these variants, we have studied p.C3575S due to its very high cosegregation with DCM and the fact that it causes strong destabilization of the parent I21 domain^19^. Our results indicate that while mutation C3575S does not affect the structure of I21, it does severely destabilize the hydrophobic core of the domain. We have then screened thousands of VUS variants for the presence of this deleterious molecular phenotype and observed that hydrophilizing variants targeting the three least solvent-accessible positions of titin Ig domains are specifically enriched in DCM cohorts. These three positions together with a strictly conserved tryptophan constitute the hydrophobic core of titin Ig domains and are fundamental for their folding^22,37^. Interestingly, this conserved tryptophan in domain I3 is the target of the highly hydrophilizing pathogenic variant *TTN* p.W976R.

Our data strongly suggest that the hydrophilizing nature of variants targeting three core residues of titin domains is an indication of pathogenicity. In **Supplementary File S1**, we provide a list with all such variants affecting the N2B isoform of titin. It is important to consider though that we have also detected this type of potentially damaging variants in a few non-DCM individuals. This is not totally unexpected if we consider that titin truncating variants, which are typically classified as pathogenic, are also found in 0.5 to 1% of the general population, at a much higher frequency than expected from the prevalence of DCM^6,14,38^. These observations highlight the need to understand downstream pathomechanisms in *TTN* related DCM^7,39,40^. In the case of the missense variants described here, it is expected that perturbation of the hydrophobic core induces strong thermodynamic destabilization of titin domains. Domain destabilization can result in protein loss of function leading to disease, as it has been observed for cardiac myosin binding protein C, a partner protein of titin in the sarcomere, in the context of hypertrophic cardiomyopathy^41^. However, in the case of a protein the size of titin, how destabilization of a single domain out of hundreds can result in such poisonous consequences is puzzling^7,16,39^. This ‘*Achilles heel*’ effect could stem from different molecular pathways, like induction of full titin degradation as observed for truncating variants^7,39^, exposure of cryptic cleavage sites altering titin mechanics^42^, or perturbed posttranslational modifications^7^, biomolecular interactions^18^ or sarcomere incorporation^16^. It is important to consider that destabilizing titin missense variants have also been linked to arrhythmogenic cardiomyopathy^43,44^ and skeletal myopathy^45^. These observations suggest that domain-specific effects downstream to domain destabilization may be important for phenotype specification.

Our work reinforces the growing evidence that the genetic landscape associating titin with DCM goes beyond truncations and includes also missense variants^19^. Traditionally, titin missense variants have been classified as VUS because, as a whole, they are not enriched in DCM cohorts^12^. However, this lack of general enrichment probably reflects the large relative weight of highly abundant, non-damaging polymorphisms. Our results illustrate that searching for specific damaging molecular features is useful to identify new DCM-causing variants. We anticipate this approach can contribute to reduce the number of DCM patients with unexplained genetic etiology and to inform development of tailored, mechanism-based preventive and therapeutic strategies^3^. Specifically, we envision that emerging high-throughput *in silico*^46–49^ and *in vitro*^50^ methods will be able to capture pathogenicity potential related to domain destabilization of all possible titin missense variants. Identification of these pathogenic variants is a prerequisite for the development of therapies able to counterbalance domain destabilization, including strategies based on exon skipping^51^ or pharmacological chaperones^52^. Furthermore, considering the broad functions played by Ig proteins^22^, which indeed are the most common family in the human genome^53^, we propose that searching for hydrophilizing variants as we have done here could benefit management of a wide range of human conditions including other (cardio)myopathies^54,55^, cancer^56^ and neurodegeneration^57^.

## Acknowledgments

JAC acknowledges funding from the Ministerio de Ciencia e Innovación (MCIN, MCIN/AEI/10.13039/501100011033) through grants BIO2017-83640-P (AEI/FEDER, EU) and PID2020-120426GB-I00, the Regional Government of Madrid (grant Tec4Bio S2018/NMT-4443, 50% co-financed by the European Social Fund and the European Regional Development Fund for the programming period 2014-2020). The CNIC is supported by the Instituto de Salud Carlos III (ISCIII), the MCIN and the Pro CNIC Foundation and is a Severo Ochoa Center of Excellence (grant CEX2020-001041-S funded by MCIN). IMM holds a fellowship from “La Caixa” Foundation (ID 100010434, fellowship code LCF/BQ/DR20/11790009) and received support from a Erasmus + Training fellowship. Financial support to DDS comes from Eusko Jaurlaritza (Basque Government) through the project IT1584-22 and from the MCIN through grants PGC2018-099321-B-I00 (AEI/FEDER, UE) and RYC-2016-19590 (AEI/FSE, EU). We thank Maria Rosaria Pricolo for analysis support with database analysis, Vytautas Gapsys for his assistance using the pmx software package, and the staff at the DIPC Supercomputing Center and the Sample Preparation and Characterization Facility at EMBL Hamburg for technical support. The synchrotron data were collected at beamline operated by EMBL Hamburg at the PETRA III storage ring (DESY, Hamburg, Germany). We thank the Spectroscopy and Nuclear Magnetic Resonance Core Unit at CNIO for access to CD instrumentation.

## METHODS

### Protein expression and purification for X-ray crystallography

Titin I21 WT cDNA was cloned in a pTEM14 expression vector using NcoI and BamHI restriction enzymes (sequence available in **Supplementary File S2**) and expressed in *E. coli* BL21 cells by overnight induction with 0.5 mM isopropyl β-D-1-thiogalactopyranoside (IPTG) at 20ºC and 250 rpm agitation. This protocol did not result in enough protein concentration in the case of the I21 C3575S domain. Hence, the mutant domain was cloned using NcoI and BamHI restriction enzymes in a pETtrx_1a plasmid vector, which includes a cleavable thioredoxin tag (TrxA) that works as a solubility enhancer for recombinant proteins expressed in *E. coli*^58^ (sequence available in **Supplementary File S2**). The protein expression protocol was similar to the one used for the WT but the temperature was decreased to 16ºC to improve protein solubility. In both cases the cells were lysed by sonication in purification buffer (30mM MES buffer pH 5.5, 100 mM NaCl, 5% glycerol for the I21 WT and 30mM sodium phosphate buffer pH 7.2, 100 mM NaCl, 5% glycerol for the I21 C3575S domain) supplemented with 20 mM imidazole, 0.01 mg/mL DNAse, 100 ug/mL lysozyme and a Complete EDTA-free protease inhibitor mixture tablet (Roche Applied Science). Then, proteins were purified using chromatography-based methods. First, proteins were purified by nickel-affinity chromatography and eluted using purification buffer including 300mM imidazole. Then, proteins were dialyzed overnight in purification buffer including 10 mM imidazole in the presence of specific proteases to remove the protein tags at a 1:100 protease:protein ratio (3C protease for the WT domain and TEV protease for the C3575S domain). After dialysis, proteins were separated from their tags by reverse nickel-affinity chromatography, concentrated and further purified by size-exclusion chromatography in purification buffer including 10 mM imidazole using the column HiLoad® 16/600 Superdex® 75 prep grade (GE healthcare). Finally, proteins were dialyzed in purification buffer to completely remove imidazole, concentrated to saturation, and stored at -80ºC. All chromatography steps were done in Fast Protein Liquid Chromatography systems (GE Healthcare).

### Protein crystallization

Crystallization conditions were screened using sitting drop vapor diffusion using 18.3 mg/mL I21 WT or 13 mg/mL I21 C3575S. To estimate concentrations, theoretical extinction coefficients were used (E^0.1%^ = 1.032, E^0.1%^ = 1.036 for WT and mutant domains, respectively)^59^. Screenings were done using 1:1 and 1:1.5 protein:mother liquor mixtures on 96-condition commercial protein crystallization screening plates. I21 WT crystallized in a buffer containing 0.06 M divalent ions (CaCl_2_, MgCl_2_), 0.1 M HEPES sodium salt, pH 7.5, 50% v/v Polyethylene glycol monomethyl ether (PEGMME) 550, PEG 20K from Morpheus HT-96 screening plate (Molecular Dimensions). I21 C3575S crystallized in 0.1 M HEPES sodium salt, pH 7.5, 25 %(w/v) PEG 2000 MME from PEGs I suite screening plate (Qiagen). Then, crystals were soaked in cryo-solution containing the crystallization mother liquid including 26% (v/v) ethylene glycol, mounted on cryo-loops (Hampton Research) and fast-cooled in liquid nitrogen.

### ray data collection and processing

Diffraction data were collected on the synchrotron radiation beamlines EMBL P14 (WT domain) and P13 (C3575S domain) at Petra III (EMBL/DESY, Hamburg) and processed using XDS^60^. I21 WT and I21 C3575S structures were solved by molecular replacement using Phaser^61^. For the I21 WT domain the reference structure was a homology model generated by I-Tasser^62^ using the 1TIT PDB structure and the I21 domain sequence. I21 C3575S structure was calculated using the solved I21 WT as a reference model. Manual modeling and refinement of both structures was carried out using COOT^63^ and Phenix^64^ (Data collection and refinement statistics in **Supplementary Table S1**). The presence of the magnesium ion in the I21 structure was confirmed using CheckMyMetal^65^. Chimera USFC^66^ was used for protein analysis and visualization. RMSD calculations refer to the structural alignment of alpha carbons in all cases. For structural comparison between WT and C3575S domains we chose chain A of the WT structure because it does not contain Mg^2+^ and chain A of the mutant, which has the best-defined electron density in the asymmetric unit.

### Protein expression and purification for biochemistry and circular dichroism

I21 WT and C3575S cDNAs were cloned in a custom made pQE80-based expression plasmid (Qiagen) using BamHI and BglII restriction enzymes (sequences available in **Supplementary File S2**). Proteins were expressed in *E. coli* BLR (DE3) in both cases. I21 WT domain was expressed by incubation with 1 mM IPTG for 3 hours at 37ºC and 250 rpm agitation. For I21 C3575S IPTG concentration was reduced to 0.4 mM and incubated at 16ºC overnight. Cells were lysed by sonication and French Press in purification buffer (50mM sodium phosphate buffer pH 7, 300 mM NaCl) supplemented with 5 ug/mL DNaseI (Roche), 5ug/mL RNAse (Sigma-Aldrich), 100 ug/mL lysozyme (Sigma-Aldrich), 10 mM MgCl_2_ and protease inhibitor cocktail III (Calbiochem). Purification of His-tagged domains was achieved by nickel-affinity using purification buffer supplemented with 1 mM DTT and gel filtration chromatography in 50mM ammonium bicarbonate pH 7 using a Superdex 200 Increase 10/300 column (GE Healthcare) following published protocols^67^. After purification the proteins were lyophilized and stored at -20ºC.

### NEM-PEGylation assay for the assessment of cysteine redox state

Lyophilized proteins were reconstituted in 20 mM sodium phosphate pH 6.5, 50 mM NaCl. 1 μg of protein was incubated with 20 mM methoxypolyethylene glycol maleimide (mPEG, Sigma-Aldrich) in the presence of 3% SDS for 10 minutes at 60ºC. Negative control samples were prepared using 10 mM N-ethymaleimide (NEM, Sigma-Aldrich) instead of mPEG. After incubation, results were analyzed by electrophoresis using 17% SDS-PAGE gels following standard protocols^68^.

### Circular Dichroism

CD spectra were collected using a Jasco J-810 spectropolarimeter. Lyophilized proteins were reconstituted in 20 mM NaPi pH 6.5, 50 mM NaCl and tested at 0.3-0.4 mg/ml protein concentration in 0.1-cm-pathlength quartz cuvettes. To estimate concentrations, theoretical extinction coefficients were used (E^0.1%^ = 0.924, E^0.1%^ = 0.925 for WT and mutant domains, respectively)^59^. To study thermal denaturation, CD signal at 215 nm was monitored as temperature increased from 25 to 85 ºC at a rate of 30ºC/h. Temperature control was achieved using a Peltier thermoelectric system. To estimate T_m_, changes in CD signal were fit to a sigmoidal function considering a two-state unfolding process using IGOR Pro (Wavemetrics).

### Molecular Dynamics (MD) simulations and alchemical free-energy calculations

MD simulations of both WT and C3575S domain were run using Amber99sb*ILND^69^ or Charmm36^70^ force fields in explicit water using the TIP3P water model^71^. Simulations were run using a hybrid structure and a hybrid topology generated using the pmx software package and webserver^72,73^ by introducing the cysteine to serine mutation in the I21 WT pdb file. 1 μs equilibrium simulations were run for the WT (λ= 0) and mutant (λ= 1) versions of the topology. Then, changes in free energy (ΔΔG) were calculated using the protocol proposed by Aldeghi *et al*.^74^. Briefly, 200 non-equilibrium 200 ps simulations were run using 200 equidistant frames from the equilibrium trajectories. During these simulations, the λ value changed from 0 to 1 (or vice versa) at a Δλ = 1x10^−5^ per simulation step. Finally, the ΔG value was calculated from these non-equilibrium trajectories using the pmx analyze Python script, which integrates over the ∂H/∂λ curves obtained from the non-equilibrium trajectories and estimates the ΔG value using Bennet’s Acceptance Ratio. This process was repeated for the unfolded state of the protein, which was approximated by a tripeptide with the cysteine (WT) or serine (C3575S) residue surrounded by two glycines^25^. The final ΔΔG (± standard deviation error) value was calculated according to:

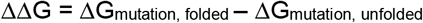

All MD simulations were run using Gromacs version 2021.4.^75^ Trajectories were analyzed using Gromacs and the MDtraj Phython library^76^. Changes in entropy associated with the variations in the distribution of rotameric states were calculated by integration of the X_1_ angle distribution according to:

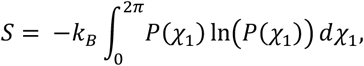

where *P*(*χ*_1_) is the probability density of the X_1_ angle and k_B_ is the Boltzmann constant^77^.

### Analysis of *TTN* variants from public databases

A database of titin missense variants was built using information from publicly available databases. The DCM-associated group was obtained from ClinVar database. We included missense variants located in Ig domains of the constitutively expressed N2B isoform of titin and showing an allele frequency (AF) < 5x10^−5^, which is compatible with conservative estimates of DCM prevalence^33,34^. The group of benign variants was populated by N2B titin missense variants from the gnomAD with AF > 5x10^−5^. For all variants, the ΔHydrophobicity value was calculated according to the Kyte-Doolittle hydrophobicity scale^31^ (ΔHydrophobicity = Hydrophobicity_mutant_ – Hydrophobicity_native_). Given the high structural similarity of titin Ig domains, SAS values were estimated according to the average SAS of equivalent positions of 13 available crystal structures of Ig domains of human titin (**Supplementary Table S2, Supplementary Figure S3**).

### Case-control study

The variants for the case-control study were obtained from the Health in Code database. The case group was built with a cohort of genotype-negative patients diagnosed with DCM who underwent genetic testing using a NGS library of 251 genes (including 121 DCM associated genes present in the Health in Code-DCM panel). The control group included patients diagnosed with diseases different from cardiomyopathies (mainly channelopathies and aortic diseases) who were referred for genetic testing and were sequenced with the same library as the case group. The odds ratio was calculated by determining the number of variants in each group that are expressed in the constitutive N2B titin, have an AF < 5x10^−5^, a SAS < 1.12 Å^2^ and that induce a ΔHydrophobicity < 2.2.

