## Supplementary Material (Figures and Tables) for "Amino acid substitutions hydrophilizing the core of titin domains cause dilated cardiomyopathy"

1 Centro Nacional de Investigaciones Cardiovasculares (CNIC), Madrid, Spain.

2 Hamburg Unit, EMBL, Hamburg, Germany.

3 Heart Failure and Inherited Cardiac Diseases Unit, Department of Cardiology, Hospital Universitario Puerta de Hierro Majadahonda, IDIPHIM, CIBERCV, Madrid, Spain.

4 Health in Code, A Coruña, Spain

5 Polimero eta Material Aurreratuak: Fisika, Kimika eta Teknologia, Kimika Fakultatea, UPV/EHU

6 Donostia International Physics Center (DIPC), Donostia-San Sebastian, Euskadi, Spain

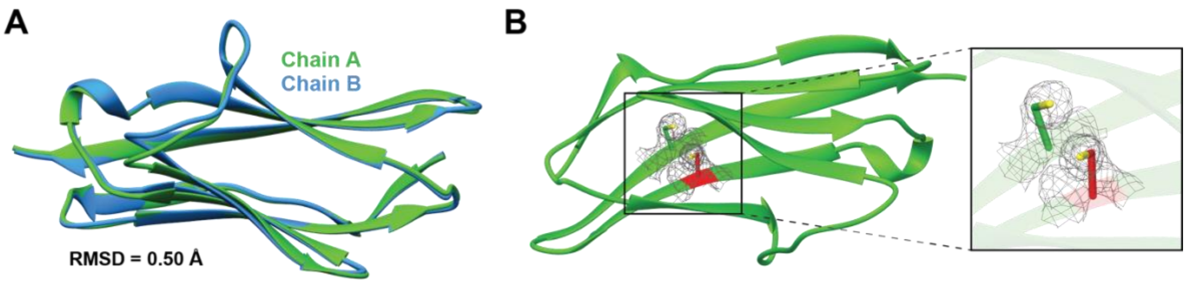


**Figure S1. Structural similarity of I21 chains and absence of disulfide bonds**: **A.** Ribbon cartoon of the structural alignment of Chain A and Chain B of I21 structure. **B.** Ribbon cartoon of I21 showing the electron density surrounding residues C3575 (red) and C3586 (green), indicating no evidence of disulfide bond formation

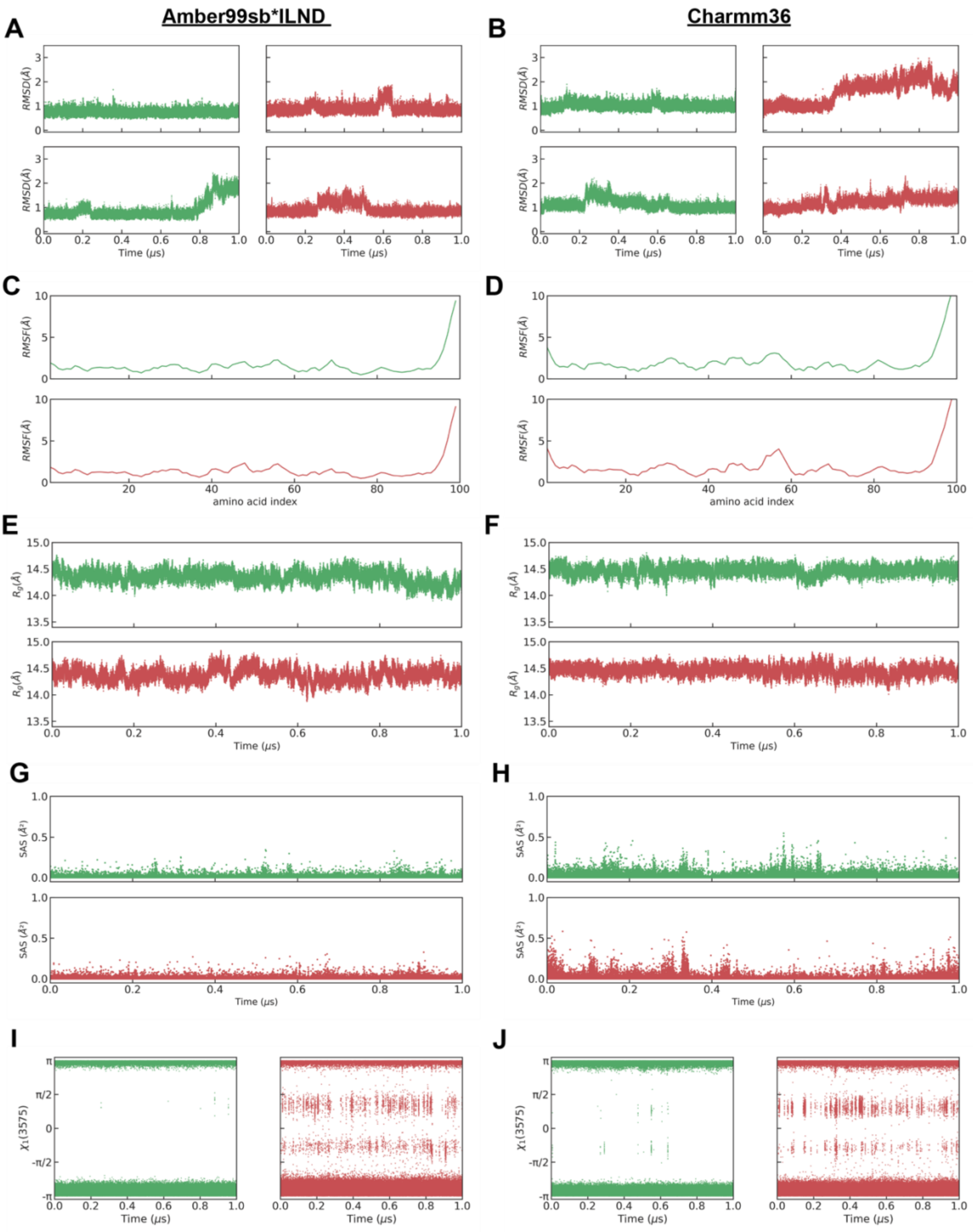


**Figure S2. Molecular dynamics simulations of the I21 WT and I21 C3575S**: **A.** Root Mean Square Displacement (RMSD) of alpha carbons over time in WT and C3575S trajectories obtained with the Amber99sb*ILND force field. **B.** RMSD of alpha carbons over time in WT and C3575S trajectories obtained with the Charmm36 force field. **C-D.** Root Mean Square Fluctuation (RMSF) of alpha carbons across protein sequence. **E-F.** Radius of gyration ($R_g$) over time in WT and C3575S trajectories. **G-H.** SAS value of position 3575 over time in WT and C3575S trajectories **I-J.** $\chi_1$ angle of position 3575 along time in WT and C3575S trajectories. Panels C-J show average data of 2 independent trajectories obtained with Amber99sb*ILND (left) or Charmm36 (right) force fields. WT data is represented in green and C3575S data in red in all panels.

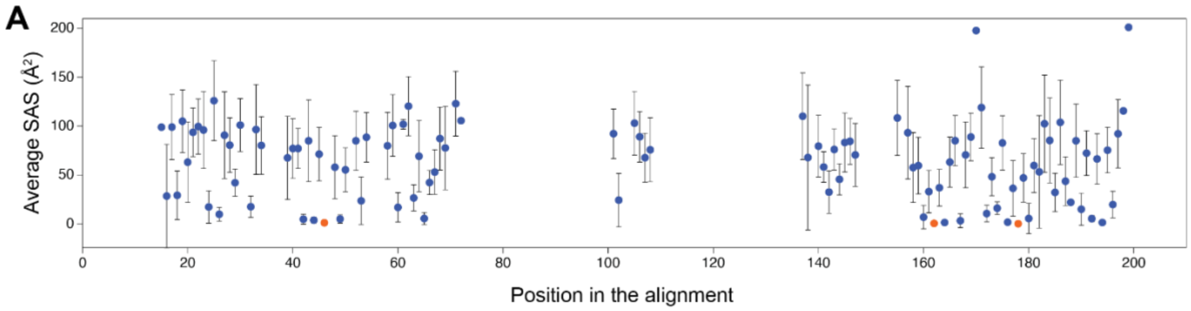

**Figure S3. Distribution of average SAS values in the alignment of titin Ig domains.** Average SAS values of titin Ig alignment positions calculated according to the SAS value of equivalent positions in 13 available crystal structures of Ig domains of human titin (**Supplementary Table S2**). Dots represent the average values, and the error bars show the standard deviation. Orange dots indicate the 3 most buried positions of titin Ig domains. Alignment positions that were not populated by any residue present in the 13 crystal structures are not represented.

### Supplementary tables

**Supplementary Table S1**. Crystallographic statistics of I21 WT (PDB ID: 8OVU) and I21 C3575S (PDB ID: 8P35): Data collection and refinement. Values in parentheses correspond to the highest resolution shell.

I21 WT (PDB ID: 8OVU)

| **Data collection** | | **Refinement** | |
|---|---|---|---|
| Radiation Source | P14 (EMBL/DESY) | $R_{work}$ (%) | 18.88 (19.43) |
| Wavelength (Å) | 0.980 | $R_{free}$ (%) | 23.89 (27.07) |
| Resolution range (Å) | 31.4 - 1.95 (2.02 - 1.95) | Number of atoms: | 1702 |
| Space group | C 2 2 21 | Macromolecules | 1512 |
| Cell dimensions: | | Ligands | 1 |
| a, b, c (Å) | 36.4 62.2 189.67 | Solvent | 189 |
| $\alpha$, $\beta$, $\gamma$ (º) | 90.0 90.0 90.0 | B factors (Å$^2$) | 33.9 |
| Unique reflections | 15822 (1041) | B $factors_{Macromolecules}$(Å$^2$) | 33.0 |
| Completeness (%) | 97.07 (91.09) | B $factors_{Ligands}$(Å$^2$) | 21.1 |
| Multiplicity | 11.3 (9.1) | B $factors_{Solvent}$(Å$^2$) | 41.1 |
| Rmerge | 0.074 (0.473) | RMSD: | |
| I/\sigma (I) | 20.2 (4.9) | Bond lengths (Å) | 0.01 |
| R-pim | 0.032 (0.228) | Bond angles (º) | 1.19 |
| CC1/2 | 0.999 (0.938) | Ramachandran plot: | |
| | | Favoured (%): | 98.97 |
| | | Allowed (%): | 1.03 |

I21 C3575S (PDB ID: 8P35)

| **Data collection** | | **Refinement** | |
|---|---|---|---|
| Radiation Source | P14 (EMBL/DESY) | $R_{work}$ (%) | 22.3 (27.1) |
| Wavelength (Å) | 0.980 | $R_{free}$ (%) | 25.91 (34.60) |

| | | | |
|---|---|---|---|
| Resolution range (Å) | 48.21 - 2.2 (2.27 - 2.20) | Number of atoms: | 4651 |
| Space group | P 21 21 21 | Macromolecules | 4471 |
| Cell dimensions: | | Ligands | 0 |
| a, b, c (Å) | 36.2 64.9 288.0 | Solvent | 180 |
| $\alpha$, $\beta$, $\gamma$ (º) | 90.0 90.0 90.0 | B factors (Å$^2$) | 42.62 |
| Unique reflections | 35852 (3088) | B factors$_{Macromolecules}$(Å$^2$) | 42.56 |
| Completeness (%) | 100 (100) | B factors$_{Ligands}$(Å$^2$) | |
| Multiplicity | 24.5 (23.6) | B factors$_{Solvent}$(Å$^2$) | 44.15 |
| Rmerge (%) | 0.085 (0.395) | RMSD: | |
| I/\sigma (I) | 22.6 (6.2) | Bond lengths (Å) | 0.014 |
| R-pim | 0.024 (0.116) | Bond angles (º) | 1.33 |
| CC1/2 | 0.995 (0.984) | Ramachandran plot: | |
| | | Favoured (%): | 99.83 |
| | | Allowed (%): | 0.17 |

**Supplementary Table S2**. PDB entries used for structure and sequence alignment with I21. The structures that contain more than one domain were split into individual domains. Domains present in more than one structure are only included once. Sequence position refers to Uniprot sequence Q8WZ42-1.

| **PDB ID** | **PDB entry name** | **Sequence position** | **Sarcomere region** | **RMSD (Å)** | **% sequence identity** |
|---|---|---|---|---|---|
| 2A38 (1st domain) | N-Terminus of titin (Z1Z2) | 1-99 | Z-disk | 0.70 | 23.2 |
| 2A38 (2nd domain) | N-Terminus of titin (Z1Z2) | 100-194 | Z-disk | 1.05 | 28.7 |
| 1G1C | I1 domain from titin | 2073-2171 | I-band | 0.63 | 24.0 |
| 5JDD (1st domain) | I9-I11 tandem from titin | 2795-2882 | I-band | 0.71 | 24.1 |
| 4QEG | I10 from titin | 2880-2967 | I-band | 0.71 | 27.3 |
| 5JDD (3rd domain) | I9-I11 tandem from titin | 2970-3053 | I-band | 0.70 | 18.4 |
| 7AHS (1st domain) | Titin-N2A Ig81-Ig83 | 9582-9672 | I-band | 0.90 | 25.6 |
| 7AHS (2nd domain) | Titin-N2A Ig81-Ig83 | 9673-9759 | I-band | 0.75 | 27.6 |
| 7AHS (3rd domain) | Titin-N2A Ig81-Ig83 | 9760-9851 | I-band | 0.84 | 28.4 |
| 1TIT | Titin, Ig repeat 27 | 12674-12765 | I-band | 1.24 | 27.9 |
| 2J8o (1st domain) | Immunoglobulin tandem repeat of titin A168-A169 | 31854-31946 | A-band | 0.67 | 30.8 |
| 2J8o (2nd domain) | Immunoglobulin tandem repeat of titin A168-A169 | 31947-32047 | A-band | 1.11 | 20.0 |
| 2BK8 | M1 domain from titin | 32497-32590 | M-line | 0.98 | 23.4 |
| 6HCI | Titin M3 domain | 32712-32861 | M-line | 0.68 | 27.8 |
| 3QP3 | Titin domain M4 | 33294-33395 | M-line | 0.77 | 20.6 |
| 1NCT | Titin module M5 | 33483-33579 | M-line | 1.28 | 22.1 |
| 3PUC | Titin domain M7 | 33774-33871 | M-line | 0.79 | 28.1 |
| 3Q5O | Human titin domain M10 | 34253-34350 | M-line | 0.88 | 26.0 |

**Supplementary Table S3:** Variants obtained in the case-control study data. Sequence position in the "Titin variant" column refer to Uniprot sequence Q8WZ42-1. Other relevant variants for selected patients are shown and annotated as pathogenic (P), likely pathogenic (LP) or variants of uncertain significance (VUS).

**Non-cardiomyopathy group**

| Case number | Titin variant | Condition | Other variants in DCM genes |
|---|---|---|---|
| 1 | M3205T | Long QT Syndrome | No other variants in DCM genes (identified LP variant in CALM2 that explains phenotype) |
| 2 | C13016G | Long QT Syndrome | No other variants in DCM genes (identified P KCNQ1 variant that explains phenotype) |
| 3 | V21725E | Brugada Syndrome | No other variants in DCM genes |
| 4 | L23925P | Long QT Syndrome | No other variants in DCM genes |

**DCM group**

| Case number | Titin variant | Condition | Other genetic variants in DCM genes |
|---|---|---|---|
| 1 | A1364T | DCM | HCN4 NP_005468.1:p.Val1150Ile NC_000015.9:g.73614986C>T (VUS) |
| 2 | A1364T | DCM | DSC2 NP_077740.1:p.Thr36Ser NC_000018.9:g.28673570T>A (VUS) |
| 3 | A1364S | DCM | No other variants in DCM genes |
| 4 | C2196Y | DCM | TTN NP_001243779.1:p.Thr30817Met NC_000002.11:g.179407110G>A (VUS) |
| 5 | C2196Y | DCM | TTN NP_001243779.1:p.Thr30817Met NC_000002.11:g.179407110G>A (VUS) |
| 6 | C3575R | DCM | No other variants in DCM genes |
| 7 | C3575S | DCM | FLNC NP_001449.3:p.Asp1816Asn NC_000007.13:g.128490904G>A (VUS) |
| 8 | C3575S | DCM | No other variants in DCM genes |
| 9 | F12838Y | DCM | ACTA1 NP_001091.1:p.Gly304Alafs*24 NC_000001.10:g.229567552delC (VUS) |
| 10 | A15628S | DCM | TTN NP_001243779.1:p.Gly31391Arg NC_000002.11:g.179403462C>G (VUS); TTN NP_001243779.1:p.Ile28818_Glu28819del NC_000002.11:g.179415878_179415883delCTCAAT (VUS) |
| 11 | A18459T | DCM | No other variants in DCM genes |
| 12 | L28271P | DCM | DSG2 NP_001934.2:p.Val276(?) NC_000018.9:g.29104548_29104550delGGT (VUS) |
| 13 | C34082G | DCM | DSC2 NM_024422.4:c.1264-4G>A NC_000018.9:g.28660322C>T (VUS) |
| 14 | C34082G | DCM | No other variants in DCM genes |
