## Supplementary File S1 for "Amino acid substitutions hydrophilizing the core of titin domains cause dilated cardiomyopathy"

List of hydrophilizing variants targeting the three core residues of titin domains and affecting the N2B isoform of titin. Sequence position refers to Uniprot sequence Q8WZ42-1.

A27R
A27N
A27D
A27E
A27Q
A27G
A27H
A27K
A27P
A27S
A27T
A27W
A27Y
L65R
L65N
L65D
L65E
L65Q
L65G
L65H
L65K
L65P
L65S
L65T
L65W
L65Y
L80R
L80N
L80D
L80E
L80Q
L80G
L80H
L80K
L80P
L80S
L80T
L80W
L80Y
V125A
V125R
V125N
V125D
V125E
V125Q
V125G
V125H
V125K
V125M
V125P

V125S
V125T
V125W
V125Y
L161R
L161N
L161D
L161E
L161Q
L161G
L161H
L161K
L161P
L161S
L161T
L161W
L161Y
V176A
V176R
V176N
V176D
V176E
V176Q
V176G
V176H
V176K
V176M
V176P
V176S
V176T
V176W
V176Y
C964R
C964N
C964D
C964E
C964Q
C964G
C964H
C964K
C964P
C964S
C964T
C964W
C964Y
L1000R
L1000N
L1000D
L1000E
L1000Q
L1000G
L1000H
L1000K
L1000P

L1000S
L1000T
L1000W
L1000Y
C1015R
C1015N
C1015D
C1015E
C1015Q
C1015G
C1015H
C1015K
C1015P
C1015S
C1015T
C1015W
C1015Y
C1103R
C1103N
C1103D
C1103E
C1103Q
C1103G
C1103H
C1103K
C1103P
C1103S
C1103T
C1103W
C1103Y
L1141R
L1141N
L1141D
L1141E
L1141Q
L1141G
L1141H
L1141K
L1141P
L1141S
L1141T
L1141W
L1141Y
I1156A
I1156R
I1156N
I1156D
I1156E
I1156Q
I1156G
I1156H
I1156K
I1156M
I1156P

I1156S
I1156T
I1156W
I1156Y
C1312R
C1312N
C1312D
C1312E
C1312Q
C1312G
C1312H
C1312K
C1312P
C1312S
C1312T
C1312W
C1312Y
L1349R
L1349N
L1349D
L1349E
L1349Q
L1349G
L1349H
L1349K
L1349P
L1349S
L1349T
L1349W
L1349Y
A1364R
A1364N
A1364D
A1364E
A1364Q
A1364G
A1364H
A1364K
A1364P
A1364S
A1364T
A1364W
A1364Y
L1478R
L1478N
L1478D
L1478E
L1478Q
L1478G
L1478H
L1478K
L1478P
L1478S
L1478T

L1478W
L1478Y
L1515R
L1515N
L1515D
L1515E
L1515Q
L1515G
L1515H
L1515K
L1515P
L1515S
L1515T
L1515W
L1515Y
V1530A
V1530R
V1530N
V1530D
V1530E
V1530Q
V1530G
V1530H
V1530K
V1530M
V1530P
V1530S
V1530T
V1530W
V1530Y
V1577A
V1577R
V1577N
V1577D
V1577E
V1577Q
V1577G
V1577H
V1577K
V1577M
V1577P
V1577S
V1577T
V1577W
V1577Y
L1615R
L1615N
L1615D
L1615E
L1615Q
L1615G
L1615H
L1615K
L1615P

L1615S
L1615T
L1615W
L1615Y
A1630R
A1630N
A1630D
A1630E
A1630Q
A1630G
A1630H
A1630K
A1630P
A1630S
A1630T
A1630W
A1630Y
C1724R
C1724N
C1724D
C1724E
C1724Q
C1724G
C1724H
C1724K
C1724P
C1724S
C1724T
C1724W
C1724Y
L1762R
L1762N
L1762D
L1762E
L1762Q
L1762G
L1762H
L1762K
L1762P
L1762S
L1762T
L1762W
L1762Y
C1777R
C1777N
C1777D
C1777E
C1777Q
C1777G
C1777H
C1777K
C1777P
C1777S
C1777T

C1777W
C1777Y
C1862R
C1862N
C1862D
C1862E
C1862Q
C1862G
C1862H
C1862K
C1862P
C1862S
C1862T
C1862W
C1862Y
L1897R
L1897N
L1897D
L1897E
L1897Q
L1897G
L1897H
L1897K
L1897P
L1897S
L1897T
L1897W
L1897Y
V1912A
V1912R
V1912N
V1912D
V1912E
V1912Q
V1912G
V1912H
V1912K
V1912M
V1912P
V1912S
V1912T
V1912W
V1912Y
V2099A
V2099R
V2099N
V2099D
V2099E
V2099Q
V2099G
V2099H
V2099K
V2099M
V2099P

V2099S
V2099T
V2099W
V2099Y
L2136R
L2136N
L2136D
L2136E
L2136Q
L2136G
L2136H
L2136K
L2136P
L2136S
L2136T
L2136W
L2136Y
V2151A
V2151R
V2151N
V2151D
V2151E
V2151Q
V2151G
V2151H
V2151K
V2151M
V2151P
V2151S
V2151T
V2151W
V2151Y
C2196R
C2196N
C2196D
C2196E
C2196Q
C2196G
C2196H
C2196K
C2196P
C2196S
C2196T
C2196W
C2196Y
L2231R
L2231N
L2231D
L2231E
L2231Q
L2231G
L2231H
L2231K
L2231P

L2231S
L2231T
L2231W
L2231Y
C2246R
C2246N
C2246D
C2246E
C2246Q
C2246G
C2246H
C2246K
C2246P
C2246S
C2246T
C2246W
C2246Y
C2288R
C2288N
C2288D
C2288E
C2288Q
C2288G
C2288H
C2288K
C2288P
C2288S
C2288T
C2288W
C2288Y
L2323R
L2323N
L2323D
L2323E
L2323Q
L2323G
L2323H
L2323K
L2323P
L2323S
L2323T
L2323W
L2323Y
F2338R
F2338N
F2338D
F2338E
F2338Q
F2338G
F2338H
F2338K
F2338P
F2338S
F2338T

F2338W
F2338Y
V2377A
V2377R
V2377N
V2377D
V2377E
V2377Q
V2377G
V2377H
V2377K
V2377M
V2377P
V2377S
V2377T
V2377W
V2377Y
L2412R
L2412N
L2412D
L2412E
L2412Q
L2412G
L2412H
L2412K
L2412P
L2412S
L2412T
L2412W
L2412Y
F2427R
F2427N
F2427D
F2427E
F2427Q
F2427G
F2427H
F2427K
F2427P
F2427S
F2427T
F2427W
F2427Y
C2466R
C2466N
C2466D
C2466E
C2466Q
C2466G
C2466H
C2466K
C2466P
C2466S
C2466T

C2466W
C2466Y
L2502R
L2502N
L2502D
L2502E
L2502Q
L2502G
L2502H
L2502K
L2502P
L2502S
L2502T
L2502W
L2502Y
L2517R
L2517N
L2517D
L2517E
L2517Q
L2517G
L2517H
L2517K
L2517P
L2517S
L2517T
L2517W
L2517Y
C2641R
C2641N
C2641D
C2641E
C2641Q
C2641G
C2641H
C2641K
C2641P
C2641S
C2641T
C2641W
C2641Y
L2676R
L2676N
L2676D
L2676E
L2676Q
L2676G
L2676H
L2676K
L2676P
L2676S
L2676T
L2676W
L2676Y

Y2691R
Y2691N
Y2691D
Y2691E
Y2691Q
Y2691K
V2728A
V2728R
V2728N
V2728D
V2728E
V2728Q
V2728G
V2728H
V2728K
V2728M
V2728P
V2728S
V2728T
V2728W
V2728Y
L2764R
L2764N
L2764D
L2764E
L2764Q
L2764G
L2764H
L2764K
L2764P
L2764S
L2764T
L2764W
L2764Y
F2779R
F2779N
F2779D
F2779E
F2779Q
F2779G
F2779H
F2779K
F2779P
F2779S
F2779T
F2779W
F2779Y
C2903R
C2903N
C2903D
C2903E
C2903Q
C2903G
C2903H

C2903K
C2903P
C2903S
C2903T
C2903W
C2903Y
L2938R
L2938N
L2938D
L2938E
L2938Q
L2938G
L2938H
L2938K
L2938P
L2938S
L2938T
L2938W
L2938Y
F2953R
F2953N
F2953D
F2953E
F2953Q
F2953G
F2953H
F2953K
F2953P
F2953S
F2953T
F2953W
F2953Y
V2990A
V2990R
V2990N
V2990D
V2990E
V2990Q
V2990G
V2990H
V2990K
V2990M
V2990P
V2990S
V2990T
V2990W
V2990Y
L3025R
L3025N
L3025D
L3025E
L3025Q
L3025G
L3025H

L3025K
L3025P
L3025S
L3025T
L3025W
L3025Y
F3040R
F3040N
F3040D
F3040E
F3040Q
F3040G
F3040H
F3040K
F3040P
F3040S
F3040T
F3040W
F3040Y
C3079R
C3079N
C3079D
C3079E
C3079Q
C3079G
C3079H
C3079K
C3079P
C3079S
C3079T
C3079W
C3079Y
L3114R
L3114N
L3114D
L3114E
L3114Q
L3114G
L3114H
L3114K
L3114P
L3114S
L3114T
L3114W
L3114Y
V3129A
V3129R
V3129N
V3129D
V3129E
V3129Q
V3129G
V3129H
V3129K

V3129M
V3129P
V3129S
V3129T
V3129W
V3129Y
F3168R
F3168N
F3168D
F3168E
F3168Q
F3168G
F3168H
F3168K
F3168P
F3168S
F3168T
F3168W
F3168Y
M3205R
M3205N
M3205D
M3205E
M3205Q
M3205G
M3205H
M3205K
M3205P
M3205S
M3205T
M3205W
M3205Y
F3220R
F3220N
F3220D
F3220E
F3220Q
F3220G
F3220H
F3220K
F3220P
F3220S
F3220T
F3220W
F3220Y
A3260R
A3260N
A3260D
A3260E
A3260Q
A3260G
A3260H
A3260K
A3260P

A3260S
A3260T
A3260W
A3260Y
L3296R
L3296N
L3296D
L3296E
L3296Q
L3296G
L3296H
L3296K
L3296P
L3296S
L3296T
L3296W
L3296Y
C3311R
C3311N
C3311D
C3311E
C3311Q
C3311G
C3311H
C3311K
C3311P
C3311S
C3311T
C3311W
C3311Y
C3366R
C3366N
C3366D
C3366E
C3366Q
C3366G
C3366H
C3366K
C3366P
C3366S
C3366T
C3366W
C3366Y
L3401R
L3401N
L3401D
L3401E
L3401Q
L3401G
L3401H
L3401K
L3401P
L3401S
L3401T

L3401W
L3401Y
F3416R
F3416N
F3416D
F3416E
F3416Q
F3416G
F3416H
F3416K
F3416P
F3416S
F3416T
F3416W
F3416Y
V3524A
V3524R
V3524N
V3524D
V3524E
V3524Q
V3524G
V3524H
V3524K
V3524M
V3524P
V3524S
V3524T
V3524W
V3524Y
L3560R
L3560N
L3560D
L3560E
L3560Q
L3560G
L3560H
L3560K
L3560P
L3560S
L3560T
L3560W
L3560Y
C3575R
C3575N
C3575D
C3575E
C3575Q
C3575G
C3575H
C3575K
C3575P
C3575S
C3575T

C3575W
C3575Y
Y3642R
Y3642N
Y3642D
Y3642E
Y3642Q
Y3642K
F3679R
F3679N
F3679D
F3679E
F3679Q
F3679G
F3679H
F3679K
F3679P
F3679S
F3679T
F3679W
F3679Y
C3694R
C3694N
C3694D
C3694E
C3694Q
C3694G
C3694H
C3694K
C3694P
C3694S
C3694T
C3694W
C3694Y
L4308R
L4308N
L4308D
L4308E
L4308Q
L4308G
L4308H
L4308K
L4308P
L4308S
L4308T
L4308W
L4308Y
L4344R
L4344N
L4344D
L4344E
L4344Q
L4344G
L4344H

L4344K
L4344P
L4344S
L4344T
L4344W
L4344Y
C4360R
C4360N
C4360D
C4360E
C4360Q
C4360G
C4360H
C4360K
C4360P
C4360S
C4360T
C4360W
C4360Y
C4404R
C4404N
C4404D
C4404E
C4404Q
C4404G
C4404H
C4404K
C4404P
C4404S
C4404T
C4404W
C4404Y
L4440R
L4440N
L4440D
L4440E
L4440Q
L4440G
L4440H
L4440K
L4440P
L4440S
L4440T
L4440W
L4440Y
C4455R
C4455N
C4455D
C4455E
C4455Q
C4455G
C4455H
C4455K
C4455P

C4455S
C4455T
C4455W
C4455Y
C12067R
C12067N
C12067D
C12067E
C12067Q
C12067G
C12067H
C12067K
C12067P
C12067S
C12067T
C12067W
C12067Y
L12102R
L12102N
L12102D
L12102E
L12102Q
L12102G
L12102H
L12102K
L12102P
L12102S
L12102T
L12102W
L12102Y
C12117R
C12117N
C12117D
C12117E
C12117Q
C12117G
C12117H
C12117K
C12117P
C12117S
C12117T
C12117W
C12117Y
C12161R
C12161N
C12161D
C12161E
C12161Q
C12161G
C12161H
C12161K
C12161P
C12161S
C12161T

C12161W
C12161Y
L12196R
L12196N
L12196D
L12196E
L12196Q
L12196G
L12196H
L12196K
L12196P
L12196S
L12196T
L12196W
L12196Y
V12211A
V12211R
V12211N
V12211D
V12211E
V12211Q
V12211G
V12211H
V12211K
V12211M
V12211P
V12211S
V12211T
V12211W
V12211Y
C12254R
C12254N
C12254D
C12254E
C12254Q
C12254G
C12254H
C12254K
C12254P
C12254S
C12254T
C12254W
C12254Y
L12291R
L12291N
L12291D
L12291E
L12291Q
L12291G
L12291H
L12291K
L12291P
L12291S
L12291T

L12291W
L12291Y
V12306A
V12306R
V12306N
V12306D
V12306E
V12306Q
V12306G
V12306H
V12306K
V12306M
V12306P
V12306S
V12306T
V12306W
V12306Y
A12345R
A12345N
A12345D
A12345E
A12345Q
A12345G
A12345H
A12345K
A12345P
A12345S
A12345T
A12345W
A12345Y
L12380R
L12380N
L12380D
L12380E
L12380Q
L12380G
L12380H
L12380K
L12380P
L12380S
L12380T
L12380W
L12380Y
Y12395R
Y12395N
Y12395D
Y12395E
Y12395Q
Y12395K
C12522R
C12522N
C12522D
C12522E
C12522Q

C12522G
C12522H
C12522K
C12522P
C12522S
C12522T
C12522W
C12522Y
L12557R
L12557N
L12557D
L12557E
L12557Q
L12557G
L12557H
L12557K
L12557P
L12557S
L12557T
L12557W
L12557Y
A12572R
A12572N
A12572D
A12572E
A12572Q
A12572G
A12572H
A12572K
A12572P
A12572S
A12572T
A12572W
A12572Y
C12611R
C12611N
C12611D
C12611E
C12611Q
C12611G
C12611H
C12611K
C12611P
C12611S
C12611T
C12611W
C12611Y
L12645R
L12645N
L12645D
L12645E
L12645Q
L12645G
L12645H

L12645K
L12645P
L12645S
L12645T
L12645W
L12645Y
C12660R
C12660N
C12660D
C12660E
C12660Q
C12660G
C12660H
C12660K
C12660P
C12660S
C12660T
C12660W
C12660Y
I12699A
I12699R
I12699N
I12699D
I12699E
I12699Q
I12699G
I12699H
I12699K
I12699M
I12699P
I12699S
I12699T
I12699W
I12699Y
L12734R
L12734N
L12734D
L12734E
L12734Q
L12734G
L12734H
L12734K
L12734P
L12734S
L12734T
L12734W
L12734Y
F12749R
F12749N
F12749D
F12749E
F12749Q
F12749G
F12749H

F12749K
F12749P
F12749S
F12749T
F12749W
F12749Y
C12788R
C12788N
C12788D
C12788E
C12788Q
C12788G
C12788H
C12788K
C12788P
C12788S
C12788T
C12788W
C12788Y
M12823R
M12823N
M12823D
M12823E
M12823Q
M12823G
M12823H
M12823K
M12823P
M12823S
M12823T
M12823W
M12823Y
F12838R
F12838N
F12838D
F12838E
F12838Q
F12838G
F12838H
F12838K
F12838P
F12838S
F12838T
F12838W
F12838Y
V12877A
V12877R
V12877N
V12877D
V12877E
V12877Q
V12877G
V12877H
V12877K

V12877M
V12877P
V12877S
V12877T
V12877W
V12877Y
I12912A
I12912R
I12912N
I12912D
I12912E
I12912Q
I12912G
I12912H
I12912K
I12912M
I12912P
I12912S
I12912T
I12912W
I12912Y
V12927A
V12927R
V12927N
V12927D
V12927E
V12927Q
V12927G
V12927H
V12927K
V12927M
V12927P
V12927S
V12927T
V12927W
V12927Y
C12966R
C12966N
C12966D
C12966E
C12966Q
C12966G
C12966H
C12966K
C12966P
C12966S
C12966T
C12966W
C12966Y
L13001R
L13001N
L13001D
L13001E
L13001Q

L13001G
L13001H
L13001K
L13001P
L13001S
L13001T
L13001W
L13001Y
C13016R
C13016N
C13016D
C13016E
C13016Q
C13016G
C13016H
C13016K
C13016P
C13016S
C13016T
C13016W
C13016Y
T13055R
T13055N
T13055D
T13055E
T13055Q
T13055H
T13055K
L13090R
L13090N
L13090D
L13090E
L13090Q
L13090G
L13090H
L13090K
L13090P
L13090S
L13090T
L13090W
L13090Y
F13105R
F13105N
F13105D
F13105E
F13105Q
F13105G
F13105H
F13105K
F13105P
F13105S
F13105T
F13105W
F13105Y

C13144R
C13144N
C13144D
C13144E
C13144Q
C13144G
C13144H
C13144K
C13144P
C13144S
C13144T
C13144W
C13144Y
L13179R
L13179N
L13179D
L13179E
L13179Q
L13179G
L13179H
L13179K
L13179P
L13179S
L13179T
L13179W
L13179Y
L13194R
L13194N
L13194D
L13194E
L13194Q
L13194G
L13194H
L13194K
L13194P
L13194S
L13194T
L13194W
L13194Y
C13233R
C13233N
C13233D
C13233E
C13233Q
C13233G
C13233H
C13233K
C13233P
C13233S
C13233T
C13233W
C13233Y
L13268R
L13268N

L13268D
L13268E
L13268Q
L13268G
L13268H
L13268K
L13268P
L13268S
L13268T
L13268W
L13268Y
C13283R
C13283N
C13283D
C13283E
C13283Q
C13283G
C13283H
C13283K
C13283P
C13283S
C13283T
C13283W
C13283Y
C13322R
C13322N
C13322D
C13322E
C13322Q
C13322G
C13322H
C13322K
C13322P
C13322S
C13322T
C13322W
C13322Y
L13357R
L13357N
L13357D
L13357E
L13357Q
L13357G
L13357H
L13357K
L13357P
L13357S
L13357T
L13357W
L13357Y
C13372R
C13372N
C13372D
C13372E

C13372Q
C13372G
C13372H
C13372K
C13372P
C13372S
C13372T
C13372W
C13372Y
C13411R
C13411N
C13411D
C13411E
C13411Q
C13411G
C13411H
C13411K
C13411P
C13411S
C13411T
C13411W
C13411Y
L13446R
L13446N
L13446D
L13446E
L13446Q
L13446G
L13446H
L13446K
L13446P
L13446S
L13446T
L13446W
L13446Y
C13461R
C13461N
C13461D
C13461E
C13461Q
C13461G
C13461H
C13461K
C13461P
C13461S
C13461T
C13461W
C13461Y
C13500R
C13500N
C13500D
C13500E
C13500Q
C13500G

C13500H
C13500K
C13500P
C13500S
C13500T
C13500W
C13500Y
L13535R
L13535N
L13535D
L13535E
L13535Q
L13535G
L13535H
L13535K
L13535P
L13535S
L13535T
L13535W
L13535Y
V13550A
V13550R
V13550N
V13550D
V13550E
V13550Q
V13550G
V13550H
V13550K
V13550M
V13550P
V13550S
V13550T
V13550W
V13550Y
C13588R
C13588N
C13588D
C13588E
C13588Q
C13588G
C13588H
C13588K
C13588P
C13588S
C13588T
C13588W
C13588Y
L13623R
L13623N
L13623D
L13623E
L13623Q
L13623G

L13623H
L13623K
L13623P
L13623S
L13623T
L13623W
L13623Y
V13638A
V13638R
V13638N
V13638D
V13638E
V13638Q
V13638G
V13638H
V13638K
V13638M
V13638P
V13638S
V13638T
V13638W
V13638Y
C13682R
C13682N
C13682D
C13682E
C13682Q
C13682G
C13682H
C13682K
C13682P
C13682S
C13682T
C13682W
C13682Y
L13717R
L13717N
L13717D
L13717E
L13717Q
L13717G
L13717H
L13717K
L13717P
L13717S
L13717T
L13717W
L13717Y
V13732A
V13732R
V13732N
V13732D
V13732E
V13732Q

V13732G
V13732H
V13732K
V13732M
V13732P
V13732S
V13732T
V13732W
V13732Y
C13771R
C13771N
C13771D
C13771E
C13771Q
C13771G
C13771H
C13771K
C13771P
C13771S
C13771T
C13771W
C13771Y
L13806R
L13806N
L13806D
L13806E
L13806Q
L13806G
L13806H
L13806K
L13806P
L13806S
L13806T
L13806W
L13806Y
C13821R
C13821N
C13821D
C13821E
C13821Q
C13821G
C13821H
C13821K
C13821P
C13821S
C13821T
C13821W
C13821Y
A13860R
A13860N
A13860D
A13860E
A13860Q
A13860G

A13860H
A13860K
A13860P
A13860S
A13860T
A13860W
A13860Y
L13895R
L13895N
L13895D
L13895E
L13895Q
L13895G
L13895H
L13895K
L13895P
L13895S
L13895T
L13895W
L13895Y
F13910R
F13910N
F13910D
F13910E
F13910Q
F13910G
F13910H
F13910K
F13910P
F13910S
F13910T
F13910W
F13910Y
V13948A
V13948R
V13948N
V13948D
V13948E
V13948Q
V13948G
V13948H
V13948K
V13948M
V13948P
V13948S
V13948T
V13948W
V13948Y
F13981R
F13981N
F13981D
F13981E
F13981Q
F13981G

F13981H
F13981K
F13981P
F13981S
F13981T
F13981W
F13981Y
I13996A
I13996R
I13996N
I13996D
I13996E
I13996Q
I13996G
I13996H
I13996K
I13996M
I13996P
I13996S
I13996T
I13996W
I13996Y
V14343A
V14343R
V14343N
V14343D
V14343E
V14343Q
V14343G
V14343H
V14343K
V14343M
V14343P
V14343S
V14343T
V14343W
V14343Y
L14379R
L14379N
L14379D
L14379E
L14379Q
L14379G
L14379H
L14379K
L14379P
L14379S
L14379T
L14379W
L14379Y
L14394R
L14394N
L14394D
L14394E

L14394Q
L14394G
L14394H
L14394K
L14394P
L14394S
L14394T
L14394W
L14394Y
A14639R
A14639N
A14639D
A14639E
A14639Q
A14639G
A14639H
A14639K
A14639P
A14639S
A14639T
A14639W
A14639Y
V14675A
V14675R
V14675N
V14675D
V14675E
V14675Q
V14675G
V14675H
V14675K
V14675M
V14675P
V14675S
V14675T
V14675W
V14675Y
I14690A
I14690R
I14690N
I14690D
I14690E
I14690Q
I14690G
I14690H
I14690K
I14690M
I14690P
I14690S
I14690T
I14690W
I14690Y
V15335A
V15335R

V15335N
V15335D
V15335E
V15335Q
V15335G
V15335H
V15335K
V15335M
V15335P
V15335S
V15335T
V15335W
V15335Y
L15371R
L15371N
L15371D
L15371E
L15371Q
L15371G
L15371H
L15371K
L15371P
L15371S
L15371T
L15371W
L15371Y
I15386A
I15386R
I15386N
I15386D
I15386E
I15386Q
I15386G
I15386H
I15386K
I15386M
I15386P
I15386S
I15386T
I15386W
I15386Y
A15628R
A15628N
A15628D
A15628E
A15628Q
A15628G
A15628H
A15628K
A15628P
A15628S
A15628T
A15628W
A15628Y

L15693R
L15693N
L15693D
L15693E
L15693Q
L15693G
L15693H
L15693K
L15693P
L15693S
L15693T
L15693W
L15693Y
I15708A
I15708R
I15708N
I15708D
I15708E
I15708Q
I15708G
I15708H
I15708K
I15708M
I15708P
I15708S
I15708T
I15708W
I15708Y
A16053R
A16053N
A16053D
A16053E
A16053Q
A16053G
A16053H
A16053K
A16053P
A16053S
A16053T
A16053W
A16053Y
L16088R
L16088N
L16088D
L16088E
L16088Q
L16088G
L16088H
L16088K
L16088P
L16088S
L16088T
L16088W
L16088Y

I16103A
I16103R
I16103N
I16103D
I16103E
I16103Q
I16103G
I16103H
I16103K
I16103M
I16103P
I16103S
I16103T
I16103W
I16103Y
A16347R
A16347N
A16347D
A16347E
A16347Q
A16347G
A16347H
A16347K
A16347P
A16347S
A16347T
A16347W
A16347Y
L16389R
L16389N
L16389D
L16389E
L16389Q
L16389G
L16389H
L16389K
L16389P
L16389S
L16389T
L16389W
L16389Y
V16404A
V16404R
V16404N
V16404D
V16404E
V16404Q
V16404G
V16404H
V16404K
V16404M
V16404P
V16404S
V16404T

V16404W
V16404Y
A16759R
A16759N
A16759D
A16759E
A16759Q
A16759G
A16759H
A16759K
A16759P
A16759S
A16759T
A16759W
A16759Y
I16803A
I16803R
I16803N
I16803D
I16803E
I16803Q
I16803G
I16803H
I16803K
I16803M
I16803P
I16803S
I16803T
I16803W
I16803Y
I16818A
I16818R
I16818N
I16818D
I16818E
I16818Q
I16818G
I16818H
I16818K
I16818M
I16818P
I16818S
I16818T
I16818W
I16818Y
A17065R
A17065N
A17065D
A17065E
A17065Q
A17065G
A17065H
A17065K
A17065P

A17065S
A17065T
A17065W
A17065Y
L17108R
L17108N
L17108D
L17108E
L17108Q
L17108G
L17108H
L17108K
L17108P
L17108S
L17108T
L17108W
L17108Y
I17123A
I17123R
I17123N
I17123D
I17123E
I17123Q
I17123G
I17123H
I17123K
I17123M
I17123P
I17123S
I17123T
I17123W
I17123Y
A17473R
A17473N
A17473D
A17473E
A17473Q
A17473G
A17473H
A17473K
A17473P
A17473S
A17473T
A17473W
A17473Y
M17507R
M17507N
M17507D
M17507E
M17507Q
M17507G
M17507H
M17507K
M17507P

M17507S
M17507T
M17507W
M17507Y
L17522R
L17522N
L17522D
L17522E
L17522Q
L17522G
L17522H
L17522K
L17522P
L17522S
L17522T
L17522W
L17522Y
G17767R
G17767N
G17767D
G17767E
G17767Q
G17767H
G17767K
L17803R
L17803N
L17803D
L17803E
L17803Q
L17803G
L17803H
L17803K
L17803P
L17803S
L17803T
L17803W
L17803Y
V17818A
V17818R
V17818N
V17818D
V17818E
V17818Q
V17818G
V17818H
V17818K
V17818M
V17818P
V17818S
V17818T
V17818W
V17818Y
A18167R
A18167N

A18167D
A18167E
A18167Q
A18167G
A18167H
A18167K
A18167P
A18167S
A18167T
A18167W
A18167Y
L18201R
L18201N
L18201D
L18201E
L18201Q
L18201G
L18201H
L18201K
L18201P
L18201S
L18201T
L18201W
L18201Y
L18216R
L18216N
L18216D
L18216E
L18216Q
L18216G
L18216H
L18216K
L18216P
L18216S
L18216T
L18216W
L18216Y
A18459R
A18459N
A18459D
A18459E
A18459Q
A18459G
A18459H
A18459K
A18459P
A18459S
A18459T
A18459W
A18459Y
L18495R
L18495N
L18495D
L18495E

L18495Q
L18495G
L18495H
L18495K
L18495P
L18495S
L18495T
L18495W
L18495Y
I18510A
I18510R
I18510N
I18510D
I18510E
I18510Q
I18510G
I18510H
I18510K
I18510M
I18510P
I18510S
I18510T
I18510W
I18510Y
A18857R
A18857N
A18857D
A18857E
A18857Q
A18857G
A18857H
A18857K
A18857P
A18857S
A18857T
A18857W
A18857Y
L18893R
L18893N
L18893D
L18893E
L18893Q
L18893G
L18893H
L18893K
L18893P
L18893S
L18893T
L18893W
L18893Y
I18908A
I18908R
I18908N
I18908D

I18908E
I18908Q
I18908G
I18908H
I18908K
I18908M
I18908P
I18908S
I18908T
I18908W
I18908Y
A19150R
A19150N
A19150D
A19150E
A19150Q
A19150G
A19150H
A19150K
A19150P
A19150S
A19150T
A19150W
A19150Y
F19188R
F19188N
F19188D
F19188E
F19188Q
F19188G
F19188H
F19188K
F19188P
F19188S
F19188T
F19188W
F19188Y
V19203A
V19203R
V19203N
V19203D
V19203E
V19203Q
V19203G
V19203H
V19203K
V19203M
V19203P
V19203S
V19203T
V19203W
V19203Y
A19555R
A19555N

A19555D
A19555E
A19555Q
A19555G
A19555H
A19555K
A19555P
A19555S
A19555T
A19555W
A19555Y
L19590R
L19590N
L19590D
L19590E
L19590Q
L19590G
L19590H
L19590K
L19590P
L19590S
L19590T
L19590W
L19590Y
L19605R
L19605N
L19605D
L19605E
L19605Q
L19605G
L19605H
L19605K
L19605P
L19605S
L19605T
L19605W
L19605Y
A19847R
A19847N
A19847D
A19847E
A19847Q
A19847G
A19847H
A19847K
A19847P
A19847S
A19847T
A19847W
A19847Y
L19883R
L19883N
L19883D
L19883E

L19883Q
L19883G
L19883H
L19883K
L19883P
L19883S
L19883T
L19883W
L19883Y
L19898R
L19898N
L19898D
L19898E
L19898Q
L19898G
L19898H
L19898K
L19898P
L19898S
L19898T
L19898W
L19898Y
A20244R
A20244N
A20244D
A20244E
A20244Q
A20244G
A20244H
A20244K
A20244P
A20244S
A20244T
A20244W
A20244Y
L20280R
L20280N
L20280D
L20280E
L20280Q
L20280G
L20280H
L20280K
L20280P
L20280S
L20280T
L20280W
L20280Y
I20295A
I20295R
I20295N
I20295D
I20295E
I20295Q

I20295G
I20295H
I20295K
I20295M
I20295P
I20295S
I20295T
I20295W
I20295Y
V20642A
V20642R
V20642N
V20642D
V20642E
V20642Q
V20642G
V20642H
V20642K
V20642M
V20642P
V20642S
V20642T
V20642W
V20642Y
L20678R
L20678N
L20678D
L20678E
L20678Q
L20678G
L20678H
L20678K
L20678P
L20678S
L20678T
L20678W
L20678Y
L20693R
L20693N
L20693D
L20693E
L20693Q
L20693G
L20693H
L20693K
L20693P
L20693S
L20693T
L20693W
L20693Y
I20931A
I20931R
I20931N
I20931D

I20931E
I20931Q
I20931G
I20931H
I20931K
I20931M
I20931P
I20931S
I20931T
I20931W
I20931Y
L20967R
L20967N
L20967D
L20967E
L20967Q
L20967G
L20967H
L20967K
L20967P
L20967S
L20967T
L20967W
L20967Y
I20982A
I20982R
I20982N
I20982D
I20982E
I20982Q
I20982G
I20982H
I20982K
I20982M
I20982P
I20982S
I20982T
I20982W
I20982Y
A21327R
A21327N
A21327D
A21327E
A21327Q
A21327G
A21327H
A21327K
A21327P
A21327S
A21327T
A21327W
A21327Y
L21364R
L21364N

L21364D
L21364E
L21364Q
L21364G
L21364H
L21364K
L21364P
L21364S
L21364T
L21364W
L21364Y
I21379A
I21379R
I21379N
I21379D
I21379E
I21379Q
I21379G
I21379H
I21379K
I21379M
I21379P
I21379S
I21379T
I21379W
I21379Y
V21725A
V21725R
V21725N
V21725D
V21725E
V21725Q
V21725G
V21725H
V21725K
V21725M
V21725P
V21725S
V21725T
V21725W
V21725Y
L21759R
L21759N
L21759D
L21759E
L21759Q
L21759G
L21759H
L21759K
L21759P
L21759S
L21759T
L21759W
L21759Y

M21774R
M21774N
M21774D
M21774E
M21774Q
M21774G
M21774H
M21774K
M21774P
M21774S
M21774T
M21774W
M21774Y
I22014A
I22014R
I22014N
I22014D
I22014E
I22014Q
I22014G
I22014H
I22014K
I22014M
I22014P
I22014S
I22014T
I22014W
I22014Y
L22050R
L22050N
L22050D
L22050E
L22050Q
L22050G
L22050H
L22050K
L22050P
L22050S
L22050T
L22050W
L22050Y
L22065R
L22065N
L22065D
L22065E
L22065Q
L22065G
L22065H
L22065K
L22065P
L22065S
L22065T
L22065W
L22065Y

A22410R
A22410N
A22410D
A22410E
A22410Q
A22410G
A22410H
A22410K
A22410P
A22410S
A22410T
A22410W
A22410Y
L22446R
L22446N
L22446D
L22446E
L22446Q
L22446G
L22446H
L22446K
L22446P
L22446S
L22446T
L22446W
L22446Y
L22461R
L22461N
L22461D
L22461E
L22461Q
L22461G
L22461H
L22461K
L22461P
L22461S
L22461T
L22461W
L22461Y
V22809A
V22809R
V22809N
V22809D
V22809E
V22809Q
V22809G
V22809H
V22809K
V22809M
V22809P
V22809S
V22809T
V22809W
V22809Y

L22843R
L22843N
L22843D
L22843E
L22843Q
L22843G
L22843H
L22843K
L22843P
L22843S
L22843T
L22843W
L22843Y
L22858R
L22858N
L22858D
L22858E
L22858Q
L22858G
L22858H
L22858K
L22858P
L22858S
L22858T
L22858W
L22858Y
V23096A
V23096R
V23096N
V23096D
V23096E
V23096Q
V23096G
V23096H
V23096K
V23096M
V23096P
V23096S
V23096T
V23096W
V23096Y
L23132R
L23132N
L23132D
L23132E
L23132Q
L23132G
L23132H
L23132K
L23132P
L23132S
L23132T
L23132W
L23132Y

V23147A
V23147R
V23147N
V23147D
V23147E
V23147Q
V23147G
V23147H
V23147K
V23147M
V23147P
V23147S
V23147T
V23147W
V23147Y
A23492R
A23492N
A23492D
A23492E
A23492Q
A23492G
A23492H
A23492K
A23492P
A23492S
A23492T
A23492W
A23492Y
L23528R
L23528N
L23528D
L23528E
L23528Q
L23528G
L23528H
L23528K
L23528P
L23528S
L23528T
L23528W
L23528Y
L23543R
L23543N
L23543D
L23543E
L23543Q
L23543G
L23543H
L23543K
L23543P
L23543S
L23543T
L23543W
L23543Y

V23891A
V23891R
V23891N
V23891D
V23891E
V23891Q
V23891G
V23891H
V23891K
V23891M
V23891P
V23891S
V23891T
V23891W
V23891Y
L23925R
L23925N
L23925D
L23925E
L23925Q
L23925G
L23925H
L23925K
L23925P
L23925S
L23925T
L23925W
L23925Y
L23940R
L23940N
L23940D
L23940E
L23940Q
L23940G
L23940H
L23940K
L23940P
L23940S
L23940T
L23940W
L23940Y
V24178A
V24178R
V24178N
V24178D
V24178E
V24178Q
V24178G
V24178H
V24178K
V24178M
V24178P
V24178S
V24178T

V24178W
V24178Y
L24214R
L24214N
L24214D
L24214E
L24214Q
L24214G
L24214H
L24214K
L24214P
L24214S
L24214T
L24214W
L24214Y
I24229A
I24229R
I24229N
I24229D
I24229E
I24229Q
I24229G
I24229H
I24229K
I24229M
I24229P
I24229S
I24229T
I24229W
I24229Y
A24574R
A24574N
A24574D
A24574E
A24574Q
A24574G
A24574H
A24574K
A24574P
A24574S
A24574T
A24574W
A24574Y
L24610R
L24610N
L24610D
L24610E
L24610Q
L24610G
L24610H
L24610K
L24610P
L24610S
L24610T

L24610W
L24610Y
L24625R
L24625N
L24625D
L24625E
L24625Q
L24625G
L24625H
L24625K
L24625P
L24625S
L24625T
L24625W
L24625Y
I24973A
I24973R
I24973N
I24973D
I24973E
I24973Q
I24973G
I24973H
I24973K
I24973M
I24973P
I24973S
I24973T
I24973W
I24973Y
L25007R
L25007N
L25007D
L25007E
L25007Q
L25007G
L25007H
L25007K
L25007P
L25007S
L25007T
L25007W
L25007Y
L25022R
L25022N
L25022D
L25022E
L25022Q
L25022G
L25022H
L25022K
L25022P
L25022S
L25022T

L25022W
L25022Y
I25260A
I25260R
I25260N
I25260D
I25260E
I25260Q
I25260G
I25260H
I25260K
I25260M
I25260P
I25260S
I25260T
I25260W
I25260Y
L25296R
L25296N
L25296D
L25296E
L25296Q
L25296G
L25296H
L25296K
L25296P
L25296S
L25296T
L25296W
L25296Y
V25311A
V25311R
V25311N
V25311D
V25311E
V25311Q
V25311G
V25311H
V25311K
V25311M
V25311P
V25311S
V25311T
V25311W
V25311Y
A25656R
A25656N
A25656D
A25656E
A25656Q
A25656G
A25656H
A25656K
A25656P

A25656S
A25656T
A25656W
A25656Y
L25693R
L25693N
L25693D
L25693E
L25693Q
L25693G
L25693H
L25693K
L25693P
L25693S
L25693T
L25693W
L25693Y
L25708R
L25708N
L25708D
L25708E
L25708Q
L25708G
L25708H
L25708K
L25708P
L25708S
L25708T
L25708W
L25708Y
V26056A
V26056R
V26056N
V26056D
V26056E
V26056Q
V26056G
V26056H
V26056K
V26056M
V26056P
V26056S
V26056T
V26056W
V26056Y
L26090R
L26090N
L26090D
L26090E
L26090Q
L26090G
L26090H
L26090K
L26090P

L26090S
L26090T
L26090W
L26090Y
L26105R
L26105N
L26105D
L26105E
L26105Q
L26105G
L26105H
L26105K
L26105P
L26105S
L26105T
L26105W
L26105Y
V26343A
V26343R
V26343N
V26343D
V26343E
V26343Q
V26343G
V26343H
V26343K
V26343M
V26343P
V26343S
V26343T
V26343W
V26343Y
L26379R
L26379N
L26379D
L26379E
L26379Q
L26379G
L26379H
L26379K
L26379P
L26379S
L26379T
L26379W
L26379Y
L26394R
L26394N
L26394D
L26394E
L26394Q
L26394G
L26394H
L26394K
L26394P

L26394S
L26394T
L26394W
L26394Y
A26738R
A26738N
A26738D
A26738E
A26738Q
A26738G
A26738H
A26738K
A26738P
A26738S
A26738T
A26738W
A26738Y
L26774R
L26774N
L26774D
L26774E
L26774Q
L26774G
L26774H
L26774K
L26774P
L26774S
L26774T
L26774W
L26774Y
L26789R
L26789N
L26789D
L26789E
L26789Q
L26789G
L26789H
L26789K
L26789P
L26789S
L26789T
L26789W
L26789Y
V27135A
V27135R
V27135N
V27135D
V27135E
V27135Q
V27135G
V27135H
V27135K
V27135M
V27135P

V27135S
V27135T
V27135W
V27135Y
L27169R
L27169N
L27169D
L27169E
L27169Q
L27169G
L27169H
L27169K
L27169P
L27169S
L27169T
L27169W
L27169Y
L27184R
L27184N
L27184D
L27184E
L27184Q
L27184G
L27184H
L27184K
L27184P
L27184S
L27184T
L27184W
L27184Y
A27821R
A27821N
A27821D
A27821E
A27821Q
A27821G
A27821H
A27821K
A27821P
A27821S
A27821T
A27821W
A27821Y
I27857A
I27857R
I27857N
I27857D
I27857E
I27857Q
I27857G
I27857H
I27857K
I27857M
I27857P

I27857S
I27857T
I27857W
I27857Y
L27872R
L27872N
L27872D
L27872E
L27872Q
L27872G
L27872H
L27872K
L27872P
L27872S
L27872T
L27872W
L27872Y
I28220A
I28220R
I28220N
I28220D
I28220E
I28220Q
I28220G
I28220H
I28220K
I28220M
I28220P
I28220S
I28220T
I28220W
I28220Y
L28256R
L28256N
L28256D
L28256E
L28256Q
L28256G
L28256H
L28256K
L28256P
L28256S
L28256T
L28256W
L28256Y
L28271R
L28271N
L28271D
L28271E
L28271Q
L28271G
L28271H
L28271K
L28271P

L28271S
L28271T
L28271W
L28271Y
L28512R
L28512N
L28512D
L28512E
L28512Q
L28512G
L28512H
L28512K
L28512P
L28512S
L28512T
L28512W
L28512Y
L28548R
L28548N
L28548D
L28548E
L28548Q
L28548G
L28548H
L28548K
L28548P
L28548S
L28548T
L28548W
L28548Y
V28563A
V28563R
V28563N
V28563D
V28563E
V28563Q
V28563G
V28563H
V28563K
V28563M
V28563P
V28563S
V28563T
V28563W
V28563Y
A28907R
A28907N
A28907D
A28907E
A28907Q
A28907G
A28907H
A28907K
A28907P

A28907S
A28907T
A28907W
A28907Y
L28943R
L28943N
L28943D
L28943E
L28943Q
L28943G
L28943H
L28943K
L28943P
L28943S
L28943T
L28943W
L28943Y
V28958A
V28958R
V28958N
V28958D
V28958E
V28958Q
V28958G
V28958H
V28958K
V28958M
V28958P
V28958S
V28958T
V28958W
V28958Y
I29306A
I29306R
I29306N
I29306D
I29306E
I29306Q
I29306G
I29306H
I29306K
I29306M
I29306P
I29306S
I29306T
I29306W
I29306Y
L29340R
L29340N
L29340D
L29340E
L29340Q
L29340G
L29340H

L29340K
L29340P
L29340S
L29340T
L29340W
L29340Y
L29355R
L29355N
L29355D
L29355E
L29355Q
L29355G
L29355H
L29355K
L29355P
L29355S
L29355T
L29355W
L29355Y
V29596A
V29596R
V29596N
V29596D
V29596E
V29596Q
V29596G
V29596H
V29596K
V29596M
V29596P
V29596S
V29596T
V29596W
V29596Y
L29632R
L29632N
L29632D
L29632E
L29632Q
L29632G
L29632H
L29632K
L29632P
L29632S
L29632T
L29632W
L29632Y
L29647R
L29647N
L29647D
L29647E
L29647Q
L29647G
L29647H

L29647K
L29647P
L29647S
L29647T
L29647W
L29647Y
A29996R
A29996N
A29996D
A29996E
A29996Q
A29996G
A29996H
A29996K
A29996P
A29996S
A29996T
A29996W
A29996Y
A30032R
A30032N
A30032D
A30032E
A30032Q
A30032G
A30032H
A30032K
A30032P
A30032S
A30032T
A30032W
A30032Y
L30047R
L30047N
L30047D
L30047E
L30047Q
L30047G
L30047H
L30047K
L30047P
L30047S
L30047T
L30047W
L30047Y
V30395A
V30395R
V30395N
V30395D
V30395E
V30395Q
V30395G
V30395H
V30395K

V30395M
V30395P
V30395S
V30395T
V30395W
V30395Y
L30429R
L30429N
L30429D
L30429E
L30429Q
L30429G
L30429H
L30429K
L30429P
L30429S
L30429T
L30429W
L30429Y
I30444A
I30444R
I30444N
I30444D
I30444E
I30444Q
I30444G
I30444H
I30444K
I30444M
I30444P
I30444S
I30444T
I30444W
I30444Y
I30687A
I30687R
I30687N
I30687D
I30687E
I30687Q
I30687G
I30687H
I30687K
I30687M
I30687P
I30687S
I30687T
I30687W
I30687Y
L30723R
L30723N
L30723D
L30723E
L30723Q

L30723G
L30723H
L30723K
L30723P
L30723S
L30723T
L30723W
L30723Y
L30738R
L30738N
L30738D
L30738E
L30738Q
L30738G
L30738H
L30738K
L30738P
L30738S
L30738T
L30738W
L30738Y
I31085A
I31085R
I31085N
I31085D
I31085E
I31085Q
I31085G
I31085H
I31085K
I31085M
I31085P
I31085S
I31085T
I31085W
I31085Y
L31119R
L31119N
L31119D
L31119E
L31119Q
L31119G
L31119H
L31119K
L31119P
L31119S
L31119T
L31119W
L31119Y
L31134R
L31134N
L31134D
L31134E
L31134Q

L31134G
L31134H
L31134K
L31134P
L31134S
L31134T
L31134W
L31134Y
C31481R
C31481N
C31481D
C31481E
C31481Q
C31481G
C31481H
C31481K
C31481P
C31481S
C31481T
C31481W
C31481Y
L31517R
L31517N
L31517D
L31517E
L31517Q
L31517G
L31517H
L31517K
L31517P
L31517S
L31517T
L31517W
L31517Y
C31532R
C31532N
C31532D
C31532E
C31532Q
C31532G
C31532H
C31532K
C31532P
C31532S
C31532T
C31532W
C31532Y
V31577A
V31577R
V31577N
V31577D
V31577E
V31577Q
V31577G

V31577H
V31577K
V31577M
V31577P
V31577S
V31577T
V31577W
V31577Y
L31613R
L31613N
L31613D
L31613E
L31613Q
L31613G
L31613H
L31613K
L31613P
L31613S
L31613T
L31613W
L31613Y
V31629A
V31629R
V31629N
V31629D
V31629E
V31629Q
V31629G
V31629H
V31629K
V31629M
V31629P
V31629S
V31629T
V31629W
V31629Y
C31876R
C31876N
C31876D
C31876E
C31876Q
C31876G
C31876H
C31876K
C31876P
C31876S
C31876T
C31876W
C31876Y
L31914R
L31914N
L31914D
L31914E
L31914Q

L31914G
L31914H
L31914K
L31914P
L31914S
L31914T
L31914W
L31914Y
V31929A
V31929R
V31929N
V31929D
V31929E
V31929Q
V31929G
V31929H
V31929K
V31929M
V31929P
V31929S
V31929T
V31929W
V31929Y
I31976A
I31976R
I31976N
I31976D
I31976E
I31976Q
I31976G
I31976H
I31976K
I31976M
I31976P
I31976S
I31976T
I31976W
I31976Y
L32012R
L32012N
L32012D
L32012E
L32012Q
L32012G
L32012H
L32012K
L32012P
L32012S
L32012T
L32012W
L32012Y
V32028A
V32028R
V32028N

V32028D
V32028E
V32028Q
V32028G
V32028H
V32028K
V32028M
V32028P
V32028S
V32028T
V32028W
V32028Y
C32516R
C32516N
C32516D
C32516E
C32516Q
C32516G
C32516H
C32516K
C32516P
C32516S
C32516T
C32516W
C32516Y
L32553R
L32553N
L32553D
L32553E
L32553Q
L32553G
L32553H
L32553K
L32553P
L32553S
L32553T
L32553W
L32553Y
C32568R
C32568N
C32568D
C32568E
C32568Q
C32568G
C32568H
C32568K
C32568P
C32568S
C32568T
C32568W
C32568Y
V32638A
V32638R
V32638N

V32638D
V32638E
V32638Q
V32638G
V32638H
V32638K
V32638M
V32638P
V32638S
V32638T
V32638W
V32638Y
L32677R
L32677N
L32677D
L32677E
L32677Q
L32677G
L32677H
L32677K
L32677P
L32677S
L32677T
L32677W
L32677Y
V32692A
V32692R
V32692N
V32692D
V32692E
V32692Q
V32692G
V32692H
V32692K
V32692M
V32692P
V32692S
V32692T
V32692W
V32692Y
I32743A
I32743R
I32743N
I32743D
I32743E
I32743Q
I32743G
I32743H
I32743K
I32743M
I32743P
I32743S
I32743T
I32743W

I32743Y
L32780R
L32780N
L32780D
L32780E
L32780Q
L32780G
L32780H
L32780K
L32780P
L32780S
L32780T
L32780W
L32780Y
V32795A
V32795R
V32795N
V32795D
V32795E
V32795Q
V32795G
V32795H
V32795K
V32795M
V32795P
V32795S
V32795T
V32795W
V32795Y
L33322R
L33322N
L33322D
L33322E
L33322Q
L33322G
L33322H
L33322K
L33322P
L33322S
L33322T
L33322W
L33322Y
L33358R
L33358N
L33358D
L33358E
L33358Q
L33358G
L33358H
L33358K
L33358P
L33358S
L33358T
L33358W

L33358Y
A33373R
A33373N
A33373D
A33373E
A33373Q
A33373G
A33373H
A33373K
A33373P
A33373S
A33373T
A33373W
A33373Y
C33509R
C33509N
C33509D
C33509E
C33509Q
C33509G
C33509H
C33509K
C33509P
C33509S
C33509T
C33509W
C33509Y
F33545R
F33545N
F33545D
F33545E
F33545Q
F33545G
F33545H
F33545K
F33545P
F33545S
F33545T
F33545W
F33545Y
V33560A
V33560R
V33560N
V33560D
V33560E
V33560Q
V33560G
V33560H
V33560K
V33560M
V33560P
V33560S
V33560T
V33560W

V33560Y
V33666A
V33666R
V33666N
V33666D
V33666E
V33666Q
V33666G
V33666H
V33666K
V33666M
V33666P
V33666S
V33666T
V33666W
V33666Y
L33703R
L33703N
L33703D
L33703E
L33703Q
L33703G
L33703H
L33703K
L33703P
L33703S
L33703T
L33703W
L33703Y
C33718R
C33718N
C33718D
C33718E
C33718Q
C33718G
C33718H
C33718K
C33718P
C33718S
C33718T
C33718W
C33718Y
V33800A
V33800R
V33800N
V33800D
V33800E
V33800Q
V33800G
V33800H
V33800K
V33800M
V33800P
V33800S

V33800T
V33800W
V33800Y
L33836R
L33836N
L33836D
L33836E
L33836Q
L33836G
L33836H
L33836K
L33836P
L33836S
L33836T
L33836W
L33836Y
C33851R
C33851N
C33851D
C33851E
C33851Q
C33851G
C33851H
C33851K
C33851P
C33851S
C33851T
C33851W
C33851Y
A33987R
A33987N
A33987D
A33987E
A33987Q
A33987G
A33987H
A33987K
A33987P
A33987S
A33987T
A33987W
A33987Y
L34021R
L34021N
L34021D
L34021E
L34021Q
L34021G
L34021H
L34021K
L34021P
L34021S
L34021T
L34021W

L34021Y
C34036R
C34036N
C34036D
C34036E
C34036Q
C34036G
C34036H
C34036K
C34036P
C34036S
C34036T
C34036W
C34036Y
C34082R
C34082N
C34082D
C34082E
C34082Q
C34082G
C34082H
C34082K
C34082P
C34082S
C34082T
C34082W
C34082Y
L34118R
L34118N
L34118D
L34118E
L34118Q
L34118G
L34118H
L34118K
L34118P
L34118S
L34118T
L34118W
L34118Y
I34133A
I34133R
I34133N
I34133D
I34133E
I34133Q
I34133G
I34133H
I34133K
I34133M
I34133P
I34133S
I34133T
I34133W

I34133Y
C34277R
C34277N
C34277D
C34277E
C34277Q
C34277G
C34277H
C34277K
C34277P
C34277S
C34277T
C34277W
C34277Y
L34315R
L34315N
L34315D
L34315E
L34315Q
L34315G
L34315H
L34315K
L34315P
L34315S
L34315T
L34315W
L34315Y
L34330R
L34330N
L34330D
L34330E
L34330Q
L34330G
L34330H
L34330K
L34330P
L34330S
L34330T
L34330W
L34330Y
