## Supplementary File S2 for "Amino acid substitutions hydrophilizing the core of titin domains cause dilated cardiomyopathy"

**Proteins for X-ray crystallography**. Sequences of the recombinant I21 WT and C3575S constructs expressed for X-ray crystallography. The codon codifying for C3575 or S3575 is highlighted in red. The cleavage site of the protease is indicated as "↓".

I21 WT DNA sequence in pTEM14 expression vector:

HisTag - 3C site – NcoI restriction site- extra nucleotides to recover the frame - **I21 WT sequence** - STOP – BamHI restriction site

ATGAAACATCACCATCACCATCACTCCGCGGGTCTGGAAGTTCTGTTCCAGGGGCCCATGGGA**G**
**GCACAGGCCCGATCTTTATCAAAGAAGTTAGCAACGCAGATATTAGCATGGGTGATGTTGCAA**
**CCCTGAGCGTTACCGTTATTGGTATTCCGAAACCGAAAATCCAGTGGTTTTTTAACGGTGTTCT**
**GCTGACCCCGAGCGCAGATTACAAATTCGTTTTTGATGGTGATGATCACAGCCTGATTATCCTG**
**TTTACCAAACTGGAAGATGAAGGCGAATATACCTGTATGGCAAGCAATGATTATGGCAAAACC**
**ATTTGTAGCGCCTACCTGAAAATCAATAGTAAGGGCGAAGGT**TAAGGATCC

I21 WT expressed protein:

MKHHHHHHSAGLEVLFQ↓GPMG**GTGPIFIKEVSNADISMGDVATLSVTVIGIPKPKIQWFFNGVLLTP**
**SADYKFVFDGDDHSLIILFTKLEDEGEYTCMASNDYGKTICSAYLKINSKGEG***

I21 C3575S DNA sequence in pETtrx_1a expression vector:

HisTag - TrxA - TEV site – NcoI restriction site- extra nucleotides to recover the frame - **I21 C3575S sequence** - STOP - BamHI

ATGAAACATCACCATCACCATCACCCCATGAGCGATAAAATTATTCACCTGACTGACGACAGTTTT
GACACGGATGTACTCAAAGCGGACGGGGCGATCCTCGTCGATTTCTGGGCAGAGTGGTGCGGT
CCGTGCAAAATGATCGCCCCGATTCTGGATGAAATCGCTGACGAATATCAGGGCAAACTGACCG
TTGCAAAACTGAACATCGATCAAAACCCTGGCACTGCGCCGAAATATGGCATCCGTGGTATCCC
GACTCTGCTGCTGTTCAAAAACGGTGAAGTGGCGGCAACCAAAGTGGGTGCACTGTCTAAAGGT
CAGTTGAAAGAGTTCCTCGACGCTAACCTGGCCGGATCTGGCAGTGGTTCTGAGAATCTTTATTT
TCAGGGCGCCATGGGA**GGCACAGGCCCGATCTTTATCAAAGAAGTTAGCAACGCAGATATTAG**
**CATGGGTGATGTTGCAACCCTGAGCGTTACCGTTATTGGTATTCCGAAACCGAAAATCCAGTG**
**GTTTTTTAACGGTGTTCTGCTGACCCCGAGCGCAGATTACAAATTCGTTTTTGATGGTGATGAT**
**CACAGCCTGATTATCCTGTTTACCAAACTGGAAGATGAAGGCGAATATACCAGTATGGCAAGC**
**AATGATTATGGCAAAACCATTTGTAGCGCCTACCTGAAAATCAATAGTAAGGGCGAAGGT**TAA
GGTACC

I21 C3575S expressed protein:

MKHHHHHHPMSDKIIHLTDDSFDTDVLKADGAILVDFWAEWCGPCKMIAPILDEIADEYQGKLTVAKL
NIDQNPGTAPKYGIRGIPTLLLFKNGEVAATKVGALSKGQLKEFLDANLAGSGSGSENLYFQ↓GAMG**G**
**TGPIFIKEVSNADISMGDVATLSVTVIGIPKPKIQWFFNGVLLTPSADYKFVFDGDDHSLIILFTKLEDE**
**GEYTSMASNDYGKTICSAYLKINSKGEG***

**Proteins for circular dichroism**. Sequences of the recombinant I21 WT and Cys3575Ser constructs expressed for circular dichroism. The codon codifying for C3575 or S3575 is highlighted in red.

I21 WT DNA sequence in pQE80B expression vector:

HisTag – BamHI restriction site- **I21 WT sequence** – BglII restriction site - STOP – KpnI restriction site

ATGAGAGGATCGCATCACCATCACCATCACGGATCC**GGCACAGGCCCGATCTTTATCAAAGAA**
**GTTAGCAACGCAGATATTAGCATGGGTGATGTTGCAACCCTGAGCGTTACCGTTATTGGTATTC**
**CGAAACCGAAAATCCAGTGGTTTTTTAACGGTGTTCTGCTGACCCCGAGCGCAGATTACAAATT**
**CGTTTTTGATGGTGATGATCACAGCCTGATTATCCTGTTTACCAAACTGGAAGATGAAGGCGAA**
**TATACCTGTATGGCAAGCAATGATTATGGCAAAACCATTTGTAGCGCCTACCTGAAAATCAATA**
**GTAAGGGCGAAGGT**AGATCTTAAGGTACC

I21 WT expressed protein

MRGSHHHHHHGS**GTGPIFIKEVSNADISMGDVATLSVTVIGIPKPKIQWFFNGVLLTPSADYKFVFDG**
**DDHSLIILFTKLEDEGEYTCMASNDYGKTICSAYLKINSKGEG**RS*

I21 C3575S DNA sequence in pQE80B expression vector:

HisTag – BamHI restriction site- **I21 WT sequence** – BglII restriction site - STOP – KpnI restriction site

ATGAGAGGATCGCATCACCATCACCATCACGGATCC**GGCACAGGCCCGATCTTTATCAAAGAA**
**GTTAGCAACGCAGATATTAGCATGGGTGATGTTGCAACCCTGAGCGTTACCGTTATTGGTATTC**
**CGAAACCGAAAATCCAGTGGTTTTTTAACGGTGTTCTGCTGACCCCGAGCGCAGATTACAAATT**
**CGTTTTTGATGGTGATGATCACAGCCTGATTATCCTGTTTACCAAACTGGAAGATGAAGGCGAA**
**TATACCAGTATGGCAAGCAATGATTATGGCAAAACCATTTGTAGCGCCTACCTGAAAATCAATA**
**GTAAGGGCGAAGGT**AGATCTTAAGGTACC

I21 C3575S expressed protein

MRGSHHHHHHGS**GTGPIFIKEVSNADISMGDVATLSVTVIGIPKPKIQWFFNGVLLTPSADYKFVFDG**
**DDHSLIILFTKLEDEGEYTSMASNDYGKTICSAYLKINSKGEG**RS*
